# Nonlinear regime transitions enable reservoir computation in activator–inhibitor cellular automata

**DOI:** 10.64898/2026.09.10.750595

**Authors:** Jurgen Riedel, Chris P. Barnes, Alexey Zaikin

## Abstract

Activator–inhibitor systems are usually studied as pattern-forming media, but the same local nonlinear interactions can also shape how information is stored and separated over time. Here we first introduce a reaction–diffusion-inspired cellular automaton as a tunable nonlinear medium with two activation families, a continuous sigmoid and a logistic–step relaxer. In each case, a single family-specific parameter changes the nonlinear response or relaxation dynamics while leaving the neighbourhood wiring fixed. This provides a controlled way to move the activator–inhibitor system between collapsed, structured, saturated, and overshooting regimes, and to relate these regimes to state-level diversity and pattern compressibility. We then use reservoir computing to test whether these tuned pattern-forming regimes support fading memory, input separability, and downstream readout learning. In this role, reservoir computing is not treated as an architecture to be optimised, but as a diagnostic of the computational properties of the untrained medium. Across parameter sweeps, stronger memory and separability are concentrated near transition regions rather than distributed uniformly across parameter space. These results identify activator–inhibitor cellular automata as interpretable unconventional reservoir substrates in which effective gain, threshold, and relaxation parameters tune the balance between pattern formation, memory, and separability. More broadly, they support the view that tissue-like or physical pattern-forming media may, in principle, shift between patterning and information-processing regimes by modulating local nonlinear dynamics.

## 1 Introduction

Modern views of biological patterning trace back to reaction–diffusion (RD) theory, where local activation with longer-range inhibition drives self-organisation into spatial structure [1–4]. RD models account for diverse phenomena, from animal markings and tissue organisation to geometry-dependent effects [5–12], and continue to be refined for growth, stochasticity, and system-specific constraints [13–17]. Network-level analyses further suggest that Turing-permissive architectures are widespread [18–23], though linear criteria can mispredict persistence or selectivity away from onset and in multistable regimes [24–29].

The broader idea of “computation” spans abstract deduction, embodiment, and universality. Aristarchus used geometric reasoning to infer celestial scales [30], the Antikythera Mechanism implemented astronomical prediction in geared bronze [31, 32], and Turing formalised symbolic computation via universal machines [33]. This historical range motivates our working view of pattern formation as computation, in which biological media can be understood as systems whose local interactions and tunable state spaces generate structured transformations of information.

As a discrete surrogate that preserves locality and parallelism, cellular automata (CA) implement activator–inhibitor logic via finite-radius convolutions while exposing independent “dials” for state resolution (alphabet), nonlinearity (gain), and memory (leak), without altering the neighbourhood wiring. Empirically, mesoscopic tissue interactions can realise CA-like updates [34]; formally, finite-neighbourhood rules define well-posed dynamical systems and can support universal computation [35–37]. CA are therefore a principled, hardware-friendly substrate for RD-inspired computation.

Reservoir computing (RC) leverages a fixed nonlinear dynamical core, the reservoir, to project inputs into a high-dimensional feature space while training only a linear readout [38–40]. Successful reservoirs exhibit fading memory and rich transient separation, properties that can arise in a range of physical substrates [41–45]. Cellular automata are attractive in this context because they combine local interactions, parallel updates, rich spatiotemporal dynamics, and simple rule-based control in a compact discrete setting [46–49].

Building on this perspective, we ask not whether a cellular automaton can serve as another feature extractor, as in previous cellular-automaton reservoir studies [46–49], but how reaction–diffusion-inspired activator–inhibitor dynamics shape state-space richness, transient memory, and input separability. This view is consistent with earlier work on computation in distributed physical media, where reaction–diffusion dynamics have been interpreted as implementing computation through local interactions and propagating spatial states [50, 51], and places the study in the context of nonlinear self-organisation and emergent computation [52, 53]. In such systems, the same local rule can give rise to collapsed, structured, saturated, or overshooting behaviour that may differ in computational capacity [54]. Rather than optimising architectures or readouts for benchmark performance, as is common in reservoir-computing studies [38, 39, 54–56], we use memory and separability tasks to probe how these behaviours emerge from local nonlinear dynamics in an unconventional computing substrate [45, 56].

To address this question, we develop an activator–inhibitor cellular automaton with two activation families, a continuous sigmoid and a logistic–step relaxer. Each family has a single control parameter that changes the nonlinear response or relaxation behaviour while leaving the underlying neighbourhood wiring fixed. This formulation separates state resolution, gain, and leak, allowing pattern formation to be examined as computation in the reservoir-computing sense [38, 39, 54]. Sweeps over activator–inhibitor radii, coupling weights, and family-specific controls map the resulting regimes and relate them to memory and separability. Emergent richness is quantified using two complementary diagnostics. *Diversity of Number of States* (DNOS) counts the number of distinct discretised state levels used after coarse-graining, whereas *Diversity of Pattern Complexity* (DPC) counts the number of distinct compression-ratio levels occupied across an ensemble. DNOS captures within-pattern alphabet breadth, while DPC captures ensemble-level variation in pattern compressibility. Finally, X-bit memory and image-classification tasks test whether the identified regimes differ in fading memory and input separability, providing a regime-level view of when activator–inhibitor pattern-forming media support useful reservoir computation and whether tissue-like media may, in principle, move between patterning and information-processing regimes by modulating local nonlinear dynamics.

We begin in subsection 2.1, where we formulate the activator–inhibitor cellular automaton from local cross-talk and define the neighbourhood terms, local balance, and activation families. We then introduce the DNOS and DPC complexity measures in subsection 2.2. The pattern-formation sweep design is described in subsection 2.3, and the resulting DNOS/DPC tuning landscapes and phase structure are analysed in subsection 2.4. Turning to reservoir computing, the X-bit memory task is introduced and evaluated in subsection 3.1, while the MNIST image-classification pipeline and results are presented in subsection 3.2. Detailed implementation and reproducibility information are collected in the Methods: subsection 5.1 for the CA pattern-formation sweeps, subsection 5.2 for the X-bit protocol, subsection 5.3 for MNIST classification, and subsection 5.4 for computational environment and reproducibility.

## 2 Cellular Automaton Model

### 2.1 From Local Cross-Talk to a Discrete Rule

With the computational view in place, we formalise a cellular automaton (CA) on a two-dimensional lattice that implements short-range activation and longer-range inhibition in the sense of Young’s local activator–inhibitor model [57]. The CA provides a discrete, rule-based analogue of an activator–inhibitor reaction–diffusion system while exposing explicit control over nonlinearity, memory, and state resolution.

#### Lattice and state variables

We consider a square lattice *L* = {0, . . ., *S*−1}^2^ with periodic boundary conditions. Each site (*i, j*) carries a continuous state variable *x_i,j_*(*t*) ∈ R representing the local activator level. States are typically initialised in [0, 1]. Under the continuous sigmoid rule they remain in (0, 1), whereas the logistic–step rule can transiently leave this interval when λ ∈/ [0, 1]. For comparability across experiments, states are normalised during downstream analysis and visualisation (Sec. 2.2).

#### Neighbourhoods and local balance

For radii *R_a_, R_i_ ∈* N and positive coupling weights W*_a_*, W*_i_* > 0, the activator and inhibitor neighbourhood sums are

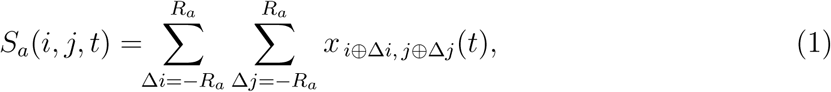

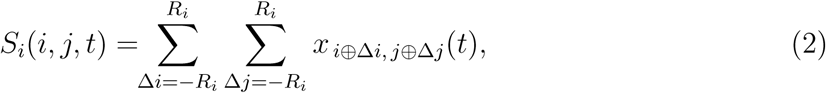

where *u ⊕ v* = (*u* + *v*) mod *S* denotes modular indexing on the torus. The local activator–inhibitor balance driving each update is then

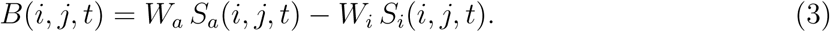

This is a discrete convolution with a compact activator kernel nested inside a broader inhibitor kernel, generating the local cross-talk that drives both self-organisation and computation.

#### Generic local update

Formally, the CA is a finite-neighbourhood, shift-commuting dynamical system. We write its local update in the generic form

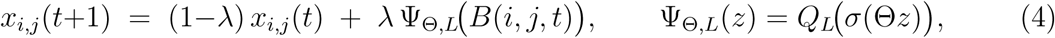

where *σ*(*u*) = (1 + *e^−u^*)*^−^*^1^ is the logistic sigmoid and *Q_L_* is an optional quantiser to *L* output levels (*Q_∞_* = id). This separates three orthogonal resources: state resolution *L*, gain Θ, and leak *λ*. In the present study we focus on the continuous case *L* = *∞* and analyse two activation families. Figure 1 summarises the continuous sigmoid rule, the smooth surrogate used for analytic comparison, and the logistic–step rule used in the discrete sweep.

**Figure 1:**
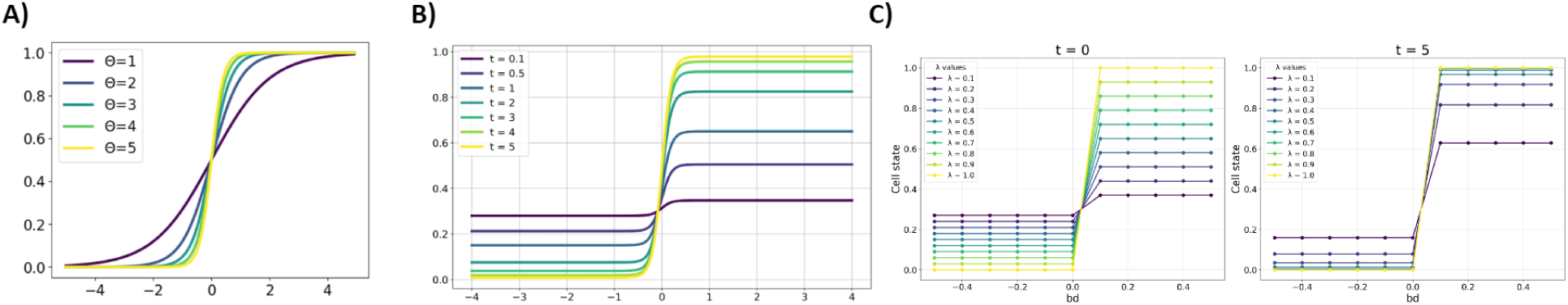
Activation families used in the CA study. All panels visualise update rules analysed in the main text, derived from the generic local update *x_t_*_+1_ = (1 − *λ*)*x_t_* + *λ* Ψ_Θ*,L*_(*B*) with Ψ_Θ*,L*_(*z*) = *Q_L_*(*σ*(Θ*z*)). **(A) Continuous sigmoid.** Transfer curves *x_t_*_+1_ = *σ*(Θ*B*) for several gains Θ, showing how increasing gain steepens the nonlinearity while the range remains (0, 1). **(B) Smooth logistic relaxation.** Continuous surrogate *f* (*B, t*; *λ,* Θ*, x*_0_) defined by Eqs. (6)–(7), shown for fractional times *t* ∈ {0.1, 0.5, 1, 2, 3, 4, 5}. The range contracts as (1 − *λ*)*^t^*, smoothly interpolating the discrete relaxation endpoints. **(C) Logistic–step (hard-target relaxation).** Update *x_t_*_+1_ = (1 − *λ*)*x_t_* + *λ H*(*B*) evaluated for *λ* ∈ [0.1, 1]. Left: initial map (*t* = 0); right: after five iterations (*t* = 5). For 0 *< λ <* 1, trajectories contract monotonically to {0, 1} with rate (1 − *λ*)*^t^*, matching the smooth limit in panel B as Θ → ∞.

#### (A) Continuous sigmoid

Setting λ = 1 and L = ∞ gives the memoryless nonlinear update

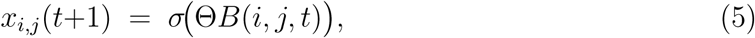

which keeps *x ∈* (0, 1). Increasing Θ steepens the transfer curve and sharpens the local decision boundary.

#### (B) Smooth logistic relaxation

For analytic comparison with the discrete rule, we introduce a smooth surrogate whose output range at effective time t matches the discrete relaxation endpoints:

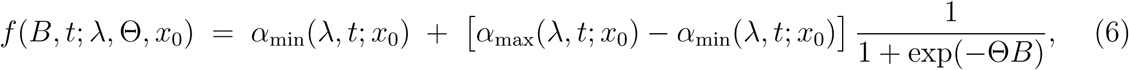

with bounds

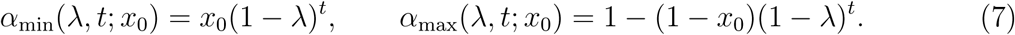

For integer *t*, this reproduces the exact lower and upper endpoints of the discrete relaxation; for fractional *t*, it provides a continuous-in-*t* envelope useful for visualisation. As *t* increases, the range contracts as (1 *− λ*)*^t^*, while larger Θ sharpens the transition near *B* = 0.

#### (C) Logistic–step (hard-target relaxation)

Taking the high-gain limit of Eq. (6) yields the Heaviside step *H*(*B*) and the hard-target update

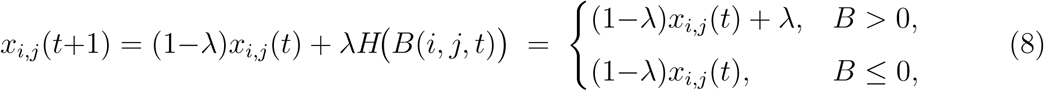

which uses only comparisons and additions. For 0 *< λ <* 1 the dynamics contract monotonically to {0, 1} with rate (1 *− λ*)*^t^*; *λ* = 1 gives an immediate threshold; *λ >* 1 produces overshoot and oscillatory convergence; and *λ <* 0 reverses the update direction.

### 2.2 Measuring Computation using Complexity Measures

To characterise the computational richness of emergent patterns, we quantify two complementary dimensions: (i) alphabet breadth, measured by the *Diversity of Number of States* (DNOS), and (ii) algorithmic depth, measured by the *Diversity of Pattern Complexity* (DPC). DNOS captures how many distinct effective state values are expressed within a single configuration, whereas DPC measures how broadly an ensemble of patterns spans the compression-ratio axis under a fixed global normalisation.

#### DNOS: precision-resolved alphabet breadth

Let the final lattice be *I ∈* ℝ*^m^*^×^*^n^*. In the current implementation, the lattice is normalised only when its values fall outside the unit interval:

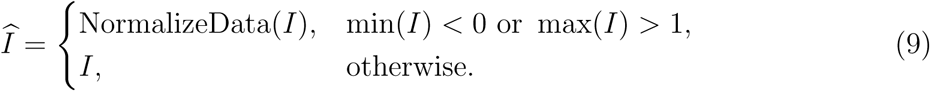

A precision-controlled representation is then formed as

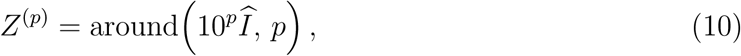

where p is the decimal-precision parameter. The Diversity of Number of States is defined as

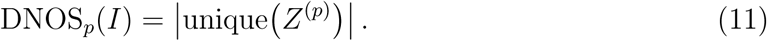

Thus, DNOS counts the number of distinct precision-resolved state values occupied on the lattice.

#### DPC: occupied compression-ratio bins across ensembles

For each image I, we compute the compression ratio

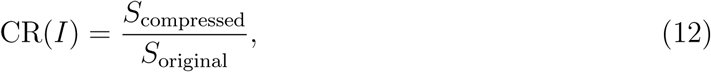

where *S*_original_ and *S*_compressed_ are the byte lengths of the original and gzip-compressed representations, respectively.

Now consider a collection of ensembles

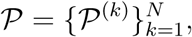

with one row of measurements associated with each ensemble. For every image in every ensemble, we evaluate its compression ratio and assemble the resulting values into a matrix **F**, with row k containing the compression ratios for ensemble *P*^(*k*)^. The full matrix is then globally normalised:

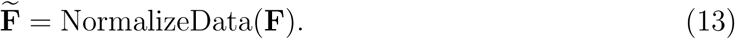

Each row is digitised into *n_b_* equally spaced bins on [0, 1]. If *h*_*j*_^(*k*)^ denotes the histogram count in bin *j* for row *k*, then the Diversity of Pattern Complexity for ensemble *k* is

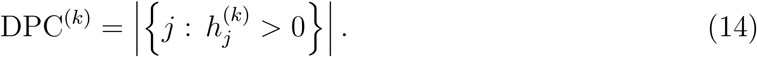

Hence, DPC measures how broadly a given ensemble occupies the globally normalised compression-ratio axis.

The two measures are therefore complementary but not symmetric. DNOS is a within-pattern count of distinct precision-resolved state values, whereas DPC is a row-wise occupied-bin count computed from globally normalised compression-ratio arrays across ensembles. In the present study these measures are not proposed as universal complexity laws; rather, they are operational coarse-grained diagnostics for detecting transitions between collapsed, structured, and overshooting nonlinear regimes.

### 2.3 Parameter sweeps with sigmoid and logistic–step activations

To characterise how activation nonlinearity shapes the dynamics of our cellular automaton (CA), we performed two systematic parameter sweeps: one using the continuous *sigmoid* activation (5) and one using its discrete analogue, the *logistic–step* rule (8). Both experiments were run on square lattices of size *S*=80 with periodic boundaries and synchronous buffered updates. The local balance followed Eq. (3), combining activation and inhibition neighbourhood sums with radii *R_a_, R_i_* and weights *W_a_, W_i_*. Initial conditions were heterogeneous, with half the sites seeded by Uniform(0, 1) draws and the remainder set to zero to promote pattern diversity.

In the *sigmoid* sweep (panel A), each site was updated using the continuous case of Eq. (5) with gain parameter Θ=*β* and *λ*=1, ensuring all states remained within (0, 1). Parameters were varied over *R_a_* ∈ {1, 2, 3, 4}, *R_i_* ∈ {2, 3, 4, 5}, *W_a_, W_i_* ∈ {0.1, 0.2, . . ., 1.0}, and *β ∈* [0.1, 30.0]. Each configuration was evolved for 60 iterations at 50% seeding. Increasing *β* sharpened the decision boundary, while varying (*R_a_, R_i_*) and the ratio *W_a_/W_i_* modulated the effective receptive field and kernel bias in the local balance (Eq. (3)).

For analytic comparison and visualisation, we also used the smooth surrogate defined by Eqs. (6)–(7) (panel C), which exactly reproduces the discrete endpoints of Eq. (8) at integer *t* and interpolates smoothly for fractional *t*. This form provides a differentiable envelope for the discrete dynamics, contracting exponentially toward {0, 1} with rate (1*−λ*)*^t^* and converging to the logistic–step limit as Θ*→∞*.

The second sweep employed the discrete hard–target relaxation (panel D), corresponding to Eq. (8). This rule contracts each site toward {0, 1} at rate (1*−λ*): for 0*<λ<*1 the convergence is monotone; *λ*=1 yields an immediate threshold; *λ>*1 introduces overshoot and oscillatory convergence; and *λ<*0 reverses the drift direction. Because the update requires only comparisons and additions, it is computationally cheaper than the sigmoid case. Parameter sweeps covered *R_a_* ∈ {1, 2, 3, 4}, *R_i_* ∈ {1, 2, 3, 4, 5}, *W_a_, W_i_* ∈ {0.1, . . ., 1.0}, and *λ ∈* [−3.0, 5.35] (with finer sampling near 0). Each configuration was evolved for 30 iterations with 50% random seeding, producing a dense grid of steady-state and transient patterns for comparison across activation regimes.

### 2.4 Complexity analysis via DNOS and DPC

For every parameter tuple in both sweeps, final lattice states were analysed with two complementary measures: DNOS (alphabet breadth) and DPC (algorithmic depth). Following Sec. 2.2, these two measures were computed by different discretisation procedures rather than by a single shared fixed-bin coarse-graining.

For DNOS, each final lattice was first checked for values outside [0, 1]; only in that case was NormalizeData applied. The lattice was then mapped to the precision-controlled representation around(10*^p^*X, p), and DNOS was taken as the number of unique values in that array (cf. Eq. (11)). DNOS therefore quantifies the number of distinct precision-resolved state values expressed within a single final pattern.

DPC was computed at the ensemble level. For each collection of final patterns associated with a fixed parameter setting, compression ratios were evaluated for all generated images, assembled into a matrix with one row per ensemble, globally normalised across all entries, digitised into n*_b_* bins on [0, 1], and reduced row-wise to the number of occupied bins (cf. Eq. (14)). Thus, DNOS measures within-pattern state-value richness, whereas DPC measures how broadly a given ensemble spans the globally normalised compression-ratio axis.

Because the two measures use different discretisation procedures, the observational scale is controlled by the DNOS precision parameter *p* and the DPC bin count *n_b_*, respectively. These settings were held fixed within each experiment to ensure comparability across parameter values.

#### Tuning the state space

We evaluated DNOS and DPC as functions of the family-specific tuning parameter, namely the sigmoid gain Θ or the logistic relaxation rate *λ*, as summarised in Figure 2. Throughout, DNOS refers to the number of unique precision-resolved state values within individual final lattices, whereas DPC reports the number of occupied bins in the corresponding globally normalised compression-ratio arrays across ensembles.

**Figure 2:**
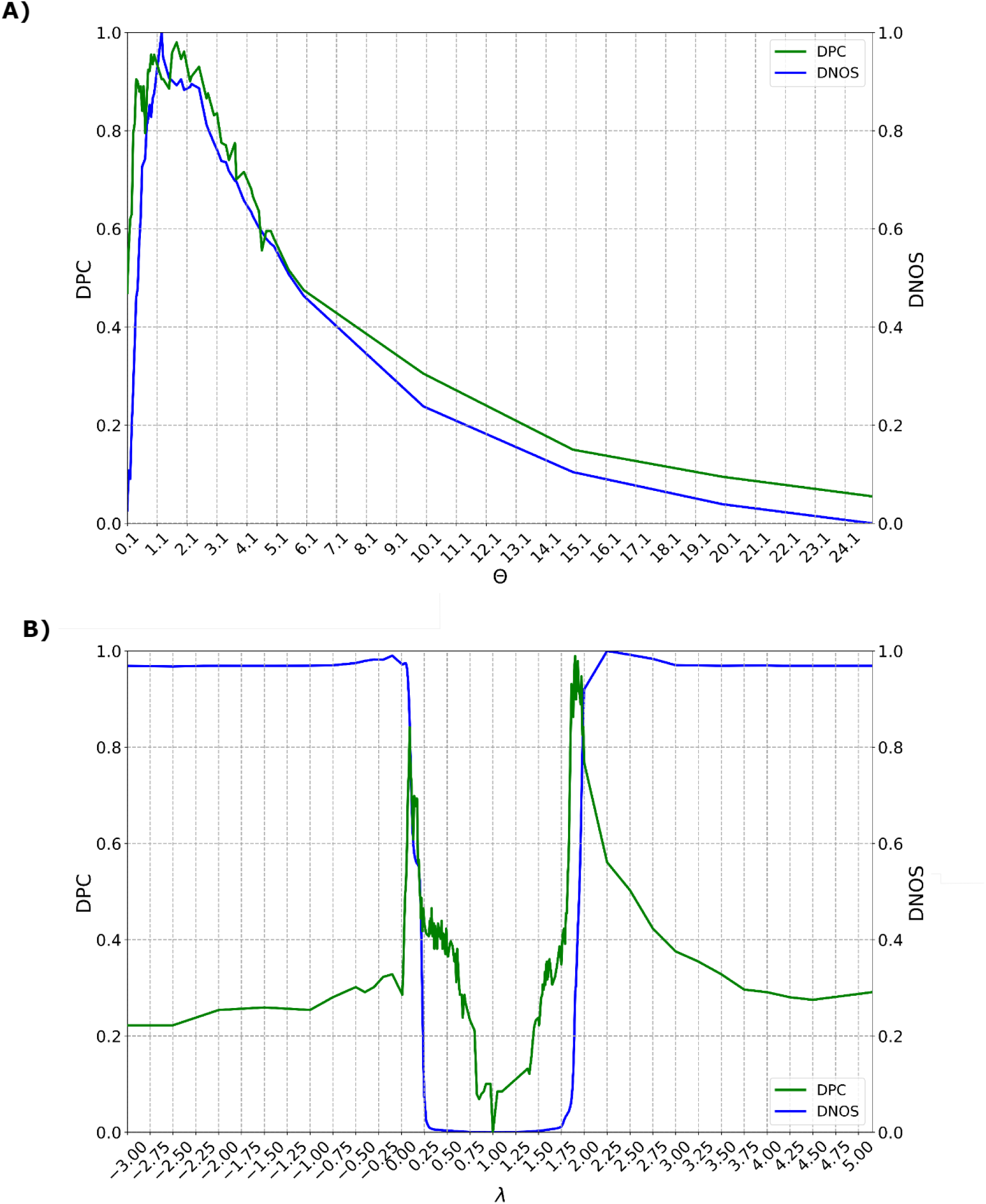
Complexity–tuning curves (composite). Panels: **(A)** sigmoid and **(B)** logistic–step activation. Each panel shows the dependence of *Diversity of Number of States* (DNOS; alphabet breadth) and *Diversity of Pattern Complexity* (DPC; algorithmic depth) on the tuning parameter, aggregated across (*R_a_, R_i_, W_a_, W_i_*) sweeps. DNOS was computed from individual final lattices as the number of unique precision-resolved state values, whereas DPC was computed across ensembles as the number of occupied bins in globally normalised compression-ratio arrays. **(A) Sigmoid:** low gain ⇒ collapse to homogeneous states (DNOS & DPC low); intermediate gain ⇒ maximal DNOS and DPC with broad multi-level usage and heterogeneous textures; high gain ⇒ saturation to near-binary domains with *frozen fronts* (DNOS remains elevated, DPC reduced). **(B) Logistic–step:** exhibits a *two-peak structure with a central minimum*: *λ <* 1 ⇒ quasi-binary relaxation (moderate DNOS, elevated DPC); *λ* = 1 ⇒ hard threshold, strictly binary after one step (DNOS drops to 1–2 occupied symbols, DPC minimal); *λ* ≈ 1.75–2.0 ⇒ overshoot induces oscillatory transients with intermediate grey levels, jointly increasing DNOS and DPC; larger *λ* ⇒ synchrony/decay, both measures decrease.

For the sigmoid rule (Figure 2A, Eq. 5), the dependence on gain Θ reveals three characteristic regimes. At low gain values (Θ ≲ 1), the response is shallow, so small neighbourhood imbalances fail to drive strong contrast formation. Patterns therefore collapse toward nearly homogeneous grey levels; DNOS remains low because only a small number of precision-resolved state values are occupied, and DPC is likewise low because the ensemble spans only a narrow range of compression-ratio bins. At intermediate gain (Θ ≈ 2–6), the sigmoid steepens enough to support broad multi-level activity and heterogeneous textures. In this regime DNOS increases because more precision-resolved state values are populated within each lattice, while DPC also reaches a maximum because the ensemble spans many distinct compression-ratio bins. This coincides with the classic transition band associated with maximal dynamical richness and computational usefulness [52, 53].

At high gain (Θ ≳ 10), the sigmoid approaches a hard threshold and interfaces between domains become pinned into stationary transition zones. These *fronts* separate regions of distinct steady states and can remain static or propagate only slowly through the lattice. Such fronts are well known in reaction–diffusion and excitable-media models as travelling or arrested activity boundaries [27, 58]. The resulting near-binary domains can still maintain non-minimal DNOS because more than one dominant state value remains occupied, but DPC drops because the ensemble becomes dominated by piecewise-constant, readily compressible patterns.

For the logistic–step rule (Figure 2B, Eq. 8), the dependence on the relaxation rate *λ* exhibits a distinct bimodal structure with two peaks separated by a central minimum. For small *λ* values (*λ ∈* [0, 0.25]), quasi-binary relaxation generates dense, irregular domain walls. DNOS remains moderate because only a limited set of precision-resolved state values is used within each lattice, but DPC shows a local maximum because the ensemble contains many structurally irregular patterns spanning distinct compression-ratio bins. At *λ* = 1, corresponding to the hard threshold, the update collapses directly to {0, 1} within a single iteration. Fronts become static, DNOS falls to the minimal binary occupancy range, and DPC also reaches a minimum because the ensemble contracts to a narrow class of efficiently compressible binary patterns.

As *λ* increases further (*λ ≈* 1.75–2.0), overshoot dynamics introduce oscillatory transients and intermediate grey levels between 0 and 1. This broadens within-pattern state usage and therefore raises DNOS, while also expanding the ensemble across multiple compression-ratio bins, producing a second DPC maximum. Such oscillatory transients and front interactions are well documented in nonlinear activator–inhibitor and excitable-media systems [27, 58]. The amplitude and width of this high-*λ* peak depend sensitively on the radii (*R_a_, R_i_*): intermediate neighbourhoods support rich mixtures of oscillatory states, very small radii suppress these oscillations, and very large radii promote synchrony across the lattice.

For *λ* values beyond this high-*λ* peak, oscillations either synchronise or decay, reducing the number of active state values within individual lattices and contracting the ensemble across fewer compression-ratio bins. Consequently, both DNOS and DPC decline. In summary, the sigmoid family displays a monotonic three-phase progression with a single maximum, whereas the logistic–step family exhibits a bimodal complexity structure with a pronounced minimum at the hard-threshold point.

#### Activation–inhibition landscapes across radii (*R_a_, R_i_*)

Figure 3 shows how *Diversity of Pattern Complexity* (DPC; A,C) and *Diversity of Number of States* (DNOS; B,D) vary over activator radius *R_a_* and inhibitor radius *R_i_* for both update families. Each tile corresponds to a fixed value of the family-specific tuning parameter: gain Θ for the sigmoid rule and relaxation rate *λ* for the logistic–step rule.

**Figure 3:**
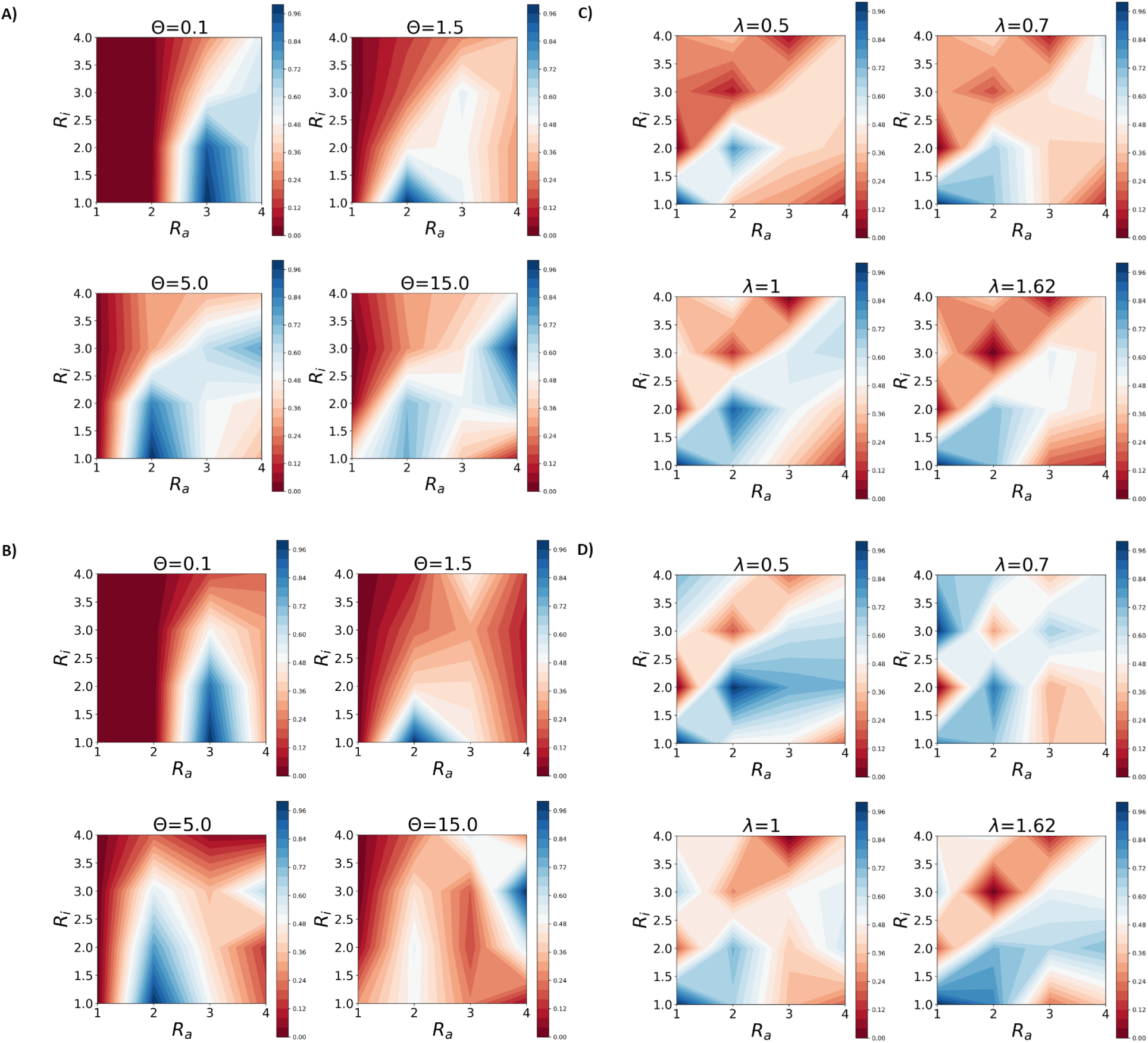
Diversity of Pattern Complexity (DPC; A,C) and Diversity of Number of States (DNOS; B,D) across activation–inhibition radii (*R_a_, R_i_*) for the sigmoid and logistic–step activation families. Each tile corresponds to a fixed value of the family-specific tuning parameter: sigmoid gain Θ in panels (A,B) and logistic–step relaxation rate *λ* in panels (C,D). **(A,B) Sigmoid:** Low gain (Θ = 0.1) yields near-homogeneous collapse and therefore low DPC/DNOS. Intermediate gain (Θ = 1.5–5.0) produces the richest regimes, with elevated DPC and broader state usage concentrated around *R_a_* ≈ 2–3, *R_i_* ≈ 2–4. At high gain (Θ = 15.0), the update approaches hard thresholding: DNOS can remain non-minimal, but DPC falls as fronts freeze into more regular, compressible domains. **(C,D) Logistic–step:** The relaxation rule shows a sharper non-monotone structure. For sub-unity *λ* (0.5–0.7), local pockets of elevated complexity/diversity appear; at *λ* = 1 the hard-threshold limit collapses the lattice to binary states and strongly reduces both measures; for *λ >* 1 (here *λ* = 1.62), overshoot reintroduces irregular high-complexity/high-diversity pockets absent in the smoother sigmoid family. DNOS captures within-pattern alphabet breadth, whereas DPC captures the breadth of occupied compression-ratio bins across ensembles, together mapping computation-friendly regions of the automaton.

Throughout this section, we use the term *fronts* to denote interfaces separating domains of distinct local states. At intermediate gain or relaxation values, such fronts remain mobile: small fluctuations along the boundary can cause them to drift, merge, or split, generating continuously evolving textures. By contrast, when the sigmoid gain becomes large, updates drive sites toward near-binary states, immobilising these interfaces and producing *frozen fronts*: piecewise-constant domain boundaries that persist with little further motion. Although frozen fronts can retain more than one active symbol (so DNOS need not be minimal), their structure is regular and readily compressible, reducing DPC at the ensemble level.

In Figure 3A, the sigmoid rule at low gain (Θ = 0.1) yields uniformly low DPC across (*R_a_, R_i_*), reflecting near-homogeneous outcomes with little structural variety, apart from a faint localised maximum near *R_a_* ≈ 3, ℝ*_i_ ≈* 2. At intermediate gain (Θ = 1.5), a broad band of elevated DPC emerges around *R_a_ ≈* 2–3, *R_i_ ≈* 2–4, consistent with a transition regime in which fronts remain mobile and irregular, generating ensembles that span many compression-complexity classes. By Θ = 5.0 this elevated region contracts toward intermediate radii as fronts begin to stabilise. At high gain (Θ = 15.0), DPC is again low over most of the plane, indicating predominantly frozen, piecewise-constant domains.

The corresponding DNOS landscapes in Figure 3B show the same broad progression with a different emphasis. At Θ = 0.1, DNOS is low throughout, with only a slight increase near *R_a_ ≈* 3, *R_i_ ≈* 2. At Θ = 1.5, modest DNOS increases appear at small *R_a_* and intermediate *R_i_*, indicating broader but still constrained state usage. At Θ = 5.0, DNOS rises more strongly around *R_a_ ≈* 2–3, *R_i_ ≈* 2–3, showing that more discretised state levels are occupied even as peripheral regions remain collapsed. By Θ = 15.0, DNOS decreases again overall, with only isolated intermediate-radius increases, consistent with near-binary frozen domains.

For the logistic–step rule, Figure 3C shows a sharper non-monotone structure. At *λ* = 0.5, a clear local DPC maximum appears near *R_a_ ≈* 2, *R_i_ ≈* 2, where contractive but irregular front dynamics generate ensembles with broad compression-ratio diversity. At *λ* = 0.7, this maximum weakens as transients damp and fronts stabilise. At *λ* = 1.0, the update becomes a strict hard threshold: the lattice binarises after a single step, domain walls become rigid, and DPC collapses nearly everywhere because the ensemble contracts to a narrow class of compressible binary patterns. At *λ* = 1.62, overshoot reintroduces dynamical activity, producing renewed local DPC peaks around *R_a_ ≈* 3, *R_i_ ≈* 2, although large regions remain simple and compressible.

The DNOS landscapes in Figure 3D parallel this behaviour. At *λ* = 0.5, DNOS peaks locally near *R_a_ ≈* 2, *R_i_ ≈* 2, indicating broader within-pattern state usage, while much of the plane remains close to quasi-binary. At *λ* = 0.7, fragmented DNOS maxima appear at small and intermediate radii, though overall breadth remains constrained. At *λ* = 1.0, DNOS collapses almost uniformly as the system reduces to the binary alphabet {0, 1} across nearly all radii. At *λ* = 1.62, overshoot generates new DNOS peaks at small *R_a_* and moderate *R_i_*, embedded within low-diversity regions where fronts synchronise.

In summary, the sigmoid family exhibits a smooth three-phase progression: low gain yields near-homogeneous collapse, intermediate gain sustains mobile fronts and maximises both DPC and DNOS, and high gain drives the system toward frozen, near-binary domains. By contrast, the logistic–step family displays a sharper bimodal structure, with richness in both a sub-unity contractive band and an overshoot-driven super-unity band, separated by a pronounced minimum at the hard-threshold point *λ* = 1. Together, these landscapes show that DNOS captures within-pattern alphabet breadth, whereas DPC captures ensemble-level diversity across compression-complexity classes, and that both identify computation-friendly regions of the automaton.

#### Pattern generation

Figure 4 A–C links global summary scores to concrete spatial codes for the sigmoid rule, while Figure 5 D–F shows the corresponding landscapes for the logistic–step rule. In both figures, the right subfigure provides exemplars for starred points: a lattice snapshot together with a *global* histogram (computed across all sites) and a grid of *local* histograms (computed from non-overlapping tiles), all using the same *B*=10^2^ coarse–graining.Histogram shape provides a visual guide to DNOS: narrow unimodal histograms indicate low DNOS (state collapse), bimodal histograms correspond to quasi-binary usage, and flat or multi-modal histograms reflect broader alphabet utilisation. Differences between global and local histograms reveal spatial heterogeneity in symbol usage. The two state-space summaries differ slightly in orientation: Figure 4 plots DPC on the x-axis and DNOS on the y-axis, whereas Figure 5 plots DNOS on the x-axis and DPC on the y-axis.

**Figure 4:**
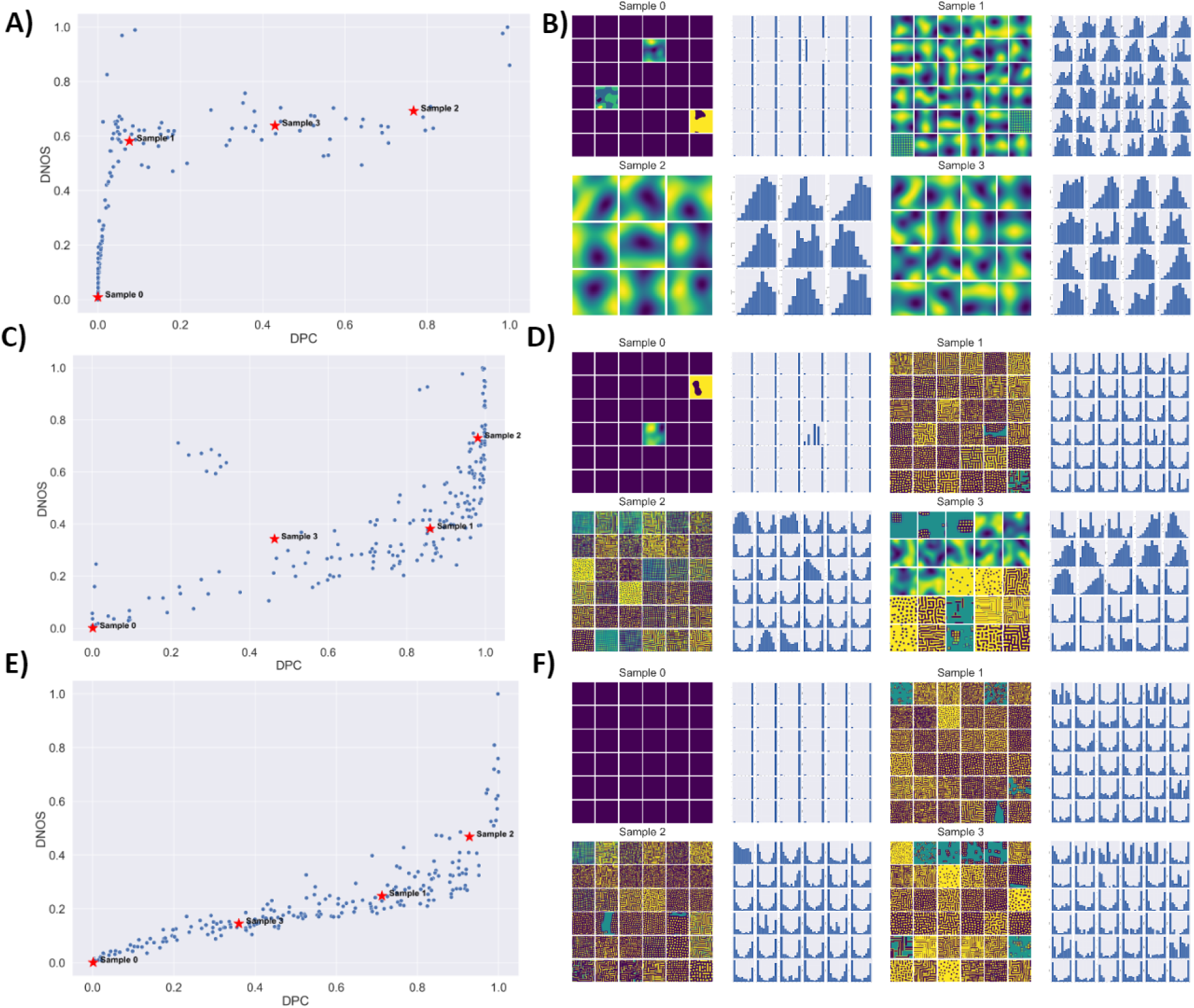
Sigmoid activation (A–C): DPC–DNOS landscapes with exemplar lattices and histograms. Left of each panel: scatter of *Diversity of Pattern Complexity* (DPC; x–axis, number of occupied compression-ratio bins across the corresponding ensemble) versus *Diversity of Number of States* (DNOS; y–axis, number of occupied bins across *B*=10^2^ uniform intervals on [0, 1]). Right of each panel: for starred points, each exemplar shows a lattice snapshot together with histogram panels (a global histogram and a grid of local histograms), all computed with the same coarse–graining (*B*=10^2^). **(A)** L-shaped cloud: DNOS rises rapidly at low DPC (“wide–but–simple”), then DPC increases while DNOS plateaus; exemplars illustrate a progression from uniform → sparse/border–biased → wide–but–simple → high-DNOS/high-DPC patterns. **(B)** Increasing gain shifts the distribution toward higher DPC while DNOS remains broadly spread; exemplars separate into *bimodal* (quasi–binary complex) and *multi–modal* (multi–level complex) regimes. **(C)** At high gain, DNOS and DPC become tightly coupled: exemplars transition from unimodal → bimodal → multi–modal histograms, reflecting the emergence of rich, multi–level textures.

**Figure 5:**
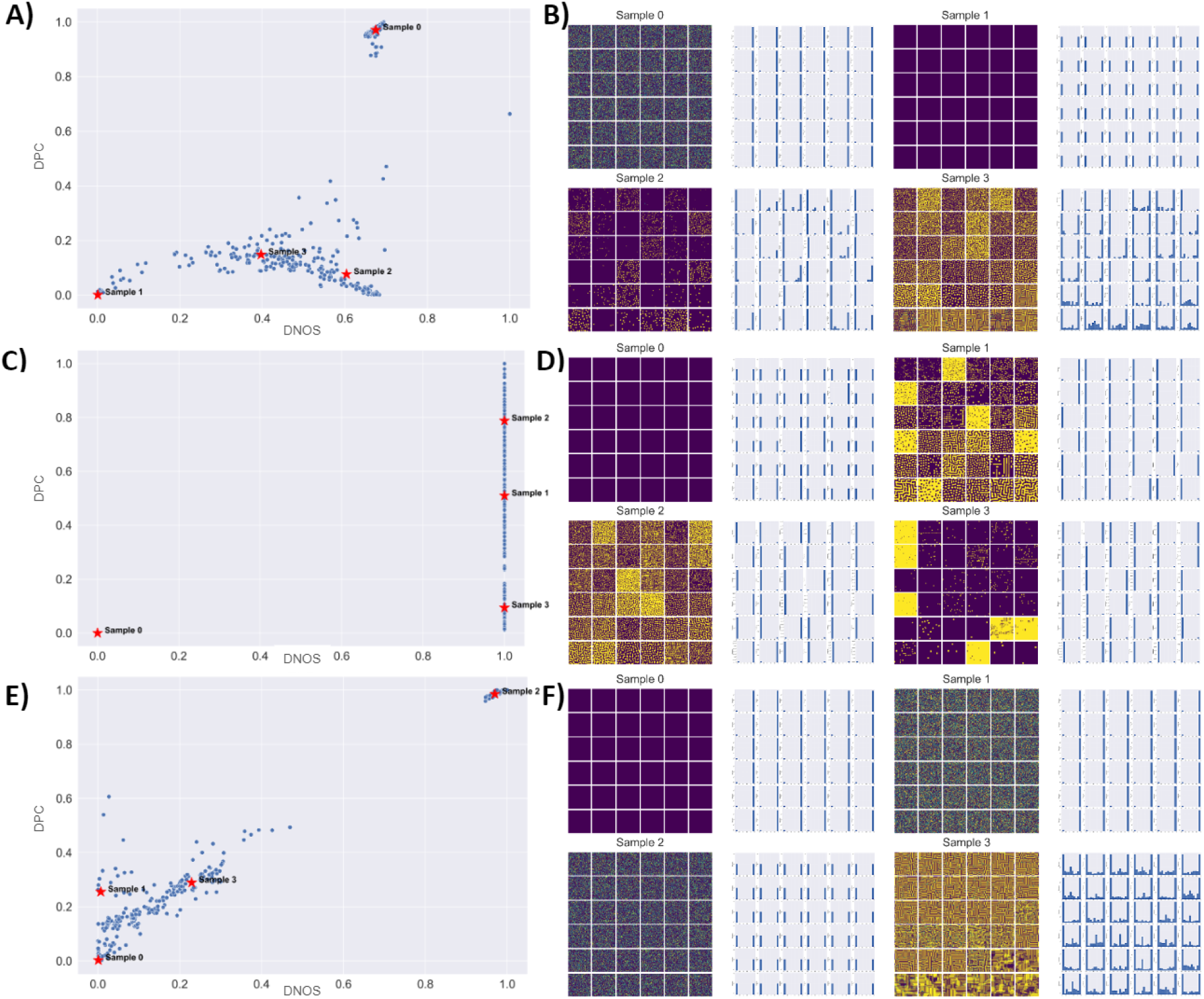
Logistic–step activation (D–F): DNOS–DPC landscapes with exemplar lattices and histograms. Same measurement protocol and histogram panels as Figure 4: each exemplar shows its lattice together with a global histogram and a grid of local histograms, all using *B*=10^2^ coarse–graining bins. In the scatter panels, DNOS is plotted on the x-axis and DPC on the y-axis. DNOS quantifies the number of occupied bins (effective alphabet size), whereas DPC quantifies the number of occupied compression-ratio bins across the corresponding ensemble. Histogram shape ↔ DNOS: narrow unimodal ↓DNOS; bimodal ≈ quasi–binary; multi–modal or flat ↑DNOS. Differences between global and local histograms reveal spatial heterogeneity in symbol usage. **(D)** *λ*=0.25 **(slow leak).** Many outcomes exhibit *bimodal* global histograms with similar local shapes (quasi–binary codes; low-to-moderate DNOS) yet moderate-to-high DPC from labyrinthine textures; a smaller pocket shows *multi–modal* histograms where mid-levels persist (higher DNOS). **(E)** *λ*=1.0 **(hard threshold).** States become strictly binary after one step; histograms collapse to two occupied bins, so DNOS reflects the relative occupancy of the two binary states, while DPC remains variable due to the spatial complexity of binary mosaics. **(F)** *λ*=1.2 **(overshoot).** Multi–level transients reappear: global and local histograms become *multi–modal* with skewed shoulders or heavy tails, indicating fluctuating symbol usage across space; DNOS–DPC trends tighten diagonally as alphabet breadth and algorithmic depth co–increase.

Across the sigmoid activation experiments (Figure 4A–C), the *DPC–DNOS* scatter reorganises systematically with gain. At baseline gain (A), the cloud forms an L–shaped envelope: DNOS rises rapidly at low DPC (“wide–but–simple” usage of many levels with repetitive spatial organisation) and then plateaus while DPC increases (incompressible textures). Exemplars illustrate the progression from uniform fields with narrow global and local histograms (low DNOS/low DPC), through sparse or border–biased patterns with weak bi-modality, to cases with broad global but repetitive local histograms (high DNOS, modest DPC), and finally multi–modal histograms with heterogeneous tiles and high DPC (the computational sweet spot). Increasing the gain (B) shifts mass toward higher DPC while preserving a wide vertical spread in DNOS; exemplars split into *bimodal* (quasi–binary yet complex) versus *multi–modal* (genuinely multi–level) regimes, evidencing a decoupling between alphabet breadth (DNOS) and spatial algorithmic depth (DPC). At high gain (C), the scatter approaches a near–monotone trend: larger DPC coincides with broader histograms (higher DNOS); exemplars move from unimodal → bimodal → multi–modal distributions, matching the joint rise in alphabet breadth and spatial complexity.

For the logistic–step rule (Figure 5D–F), the summary is shown as a *DNOS–DPC* scatter, with DNOS on the x-axis and DPC on the y-axis. Tuning *λ* induces sharper phase shifts. At *λ* = 0.25 (D), most runs are quasi–binary, clustering at low-to-moderate DNOS with moderate DPC, while a smaller subregion sustains persistent multi–level states with higher DNOS but still moderate algorithmic depth before full relaxation. At *λ* = 1 (E), the update becomes a hard threshold: lattices are strictly binary after one step, so DNOS collapses toward the binary limit, while DPC remains variable because binary mosaics can still differ substantially in spatial complexity. At *λ* = 1.2 (F), overshoot tightens the DNOS–DPC relation along an approximately diagonal trend; exemplar histograms reveal intermittency (patchy, multi–modal tiles) and, at peaks, jointly high DNOS and high DPC.

## 3 Reservoir computing with activator–inhibitor cellular automata

The CA reservoir-computing framework proceeds in three stages [46–48]. First, the input signal **u**(*t*) is embedded in the lattice by encoding it either into the initial configuration or into time-varying boundary conditions. Second, the CA evolves for *I* discrete iterations under local update rules

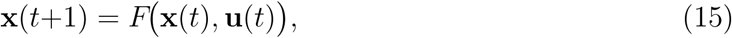

thereby projecting the data into a high-dimensional, nonlinear state space [49]. Third, a linear read-out layer maps the reservoir state to the task output via

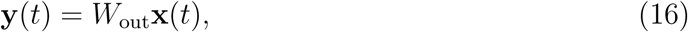

as in standard CA-based RC pipelines [46].

In one-dimensional reservoirs, elementary CA rules act on an expanded bit string whose buffer cells suppress edge artefacts; after I steps the state is

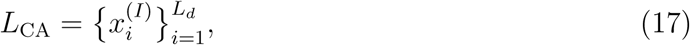

with each cell x^(*I*)^ encoding a nonlinear summary of the entire input history [48]. Two-dimensional reservoirs generalise this mechanism by mapping the signal to a lattice whose local interactions generate richer spatio-temporal transients; the snapshots collected at each iteration are flattened and concatenated,

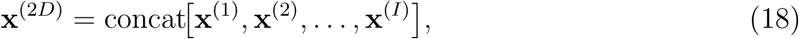

to form a feature vector compatible with the same linear read-out [49].

### 3.1 X-bit memory task

In the following sections, temporal signals are injected into the CA lattice by writing each input token into a randomly permuted subset of lattice sites at successive task steps. After each write, the lattice evolves autonomously for I reservoir updates, and a linear readout

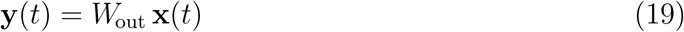

is trained to reconstruct the delayed target in the X-bit memory task, following established CA-based reservoir-computing benchmarks [46, 47].

This protocol exploits the separation between rich but fixed intrinsic dynamics and lightweight supervised learning that makes reservoir computing attractive, while simultaneously probing the computational limits of a model that is, at its core, a minimal discretised morphogenetic network. By this we mean a distributed system of local interactions that self-organises into spatial patterns: in biology, activator–inhibitor dynamics and gene-regulatory topology can generate tissue-level organisation [1, 2, 6, 19, 20]. At a mesoscopic level, CA-like update behaviour has also been observed in living tissue [34]. In the discrete case, a cellular automaton lattice realises the same organising principle through finite-radius update rules [35–37]. Framing the CA reservoir as a morphogenetic network therefore highlights a mechanistic parallel between biological pattern formation and distributed computation.

Because the update rule is local, nonlinear, and parameter-tunable, the CA acts as a fixed, high-dimensional reservoir into which external signals can be embedded through controlled perturbations of the lattice state. By demonstrating that a biologically motivated CA exhibits fading-memory behaviour and supports linearly decodable recall [48, 49], we offer both a new RC substrate and a fresh lens on how living tissues might, in principle, harness pattern-forming dynamics for distributed computation.

Reservoir memory is quantified with the X-bit task, in which the input stream is split across four channels: signal, flipped signal, distractor, and cue. The task therefore requires the system to preserve the relevant bits across a distraction period and to release them only when the cue appears. Upon cue presentation, the readout produces **y**(t), and its bit-wise recall contributes to the weighted score

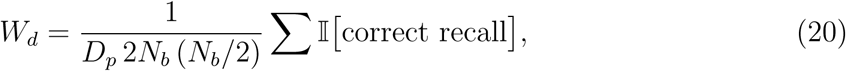

where *D_p_* is the distractor length and *N_b_* is the input size [47, 48].

In the sweeps below, we refer to four input–cue source geometries, denoted 20/30, 60/30, 20/60, and 60/60. These labels identify the four spatial write configurations used to inject the signal and cue streams on the lattice, allowing us to test how sensitive memory performance is to the geometry of the write process while keeping the reservoir dynamics and readout protocol otherwise fixed.

Parameter sweeps in the CA–RC literature show that performance generally improves with the iteration count *I* and with the effective feature-space expansion induced by the reservoir, albeit at increased computational cost; the reservoir side length *L_d_* likewise modulates accuracy [47, 48]. Across tasks with pronounced spatio-temporal structure, two-dimensional reservoirs can outperform one-dimensional ones because their local interactions generate richer transients [48, 49]. Consistent with earlier CA-reservoir studies, regimes that balance stability with transient separability appear more supportive of recall than regimes that collapse rapidly to homogeneous or rigidly binary states [48, 49].

The computational capacity of a cellular-automaton reservoir is constrained in part by the cardinality of its effective state alphabet: an N-cell lattice with |S| distinct symbols spans a state space of size |S|*^N^*, which grows exponentially with system size. In RC terms, richer effective alphabets can enlarge the accessible feature space [49].

A convenient parametrisation of the reservoir dynamics employs the two activation families introduced earlier: the continuous sigmoid rule (Eq. 5) and the logistic–step relaxation rule (Eq. 8). These families do not share the same tuning variable. For the sigmoid rule, the control parameter is the gain Θ, which sets the steepness of the nonlinear response. For the logistic–step rule, the control parameter is the relaxation rate *λ*, which governs contraction, hard-thresholding at *λ* = 1, and overshoot for *λ >* 1. Low Θ yields a shallow, near-linear sigmoid, whereas high Θ approaches a hard threshold; by contrast, low *λ* yields gradual relaxation and high *λ* can induce oscillatory or overshooting behaviour. Together, these two families provide a controlled way to move between graded and near-discrete reservoir dynamics without changing the underlying local wiring. In our experiments, the best memory performance is obtained not in fully collapsed or rigidly saturated regimes, but in intermediate operating regions that preserve both fading memory and nonlinear separability [48, 49].

Extensive experiments confirm that carefully tuned cellular automata can serve as effective reservoirs for temporal-memory tasks, especially when operated in regimes that preserve rich but decodable spatiotemporal transients rather than collapsing immediately to homogeneous or rigidly binary states [48, 49, 59]. Together, these results reinforce the view that CA provide a compact, efficient, and potentially hardware-friendly substrate for reservoir computing.

Figure 6 summarises the X-bit memory score W*_d_* as a function of the family-specific tuning parameter for the four input–cue source geometries described in Sec. 3.1, under the protocol of Methods Sec. 6.2. For the sigmoid family (Figure 6A), performance varies systematically with both source geometry and gain Θ: all four curves show a broad geometry-dependent operating range, with a shallow depression at intermediate gain and partial recovery at larger Θ. For the logistic–step family (Figure 6B), the dependence is sharper: all four source geometries exhibit a pronounced minimum near the hard-threshold regime *λ ≈* 0.8–1.0, followed by recovery away from this region. Thus, the threshold-like regime is structurally unfavourable on average for the X-bit task, whereas both more contractive and mildly overshooting settings recover stronger recall.

**Figure 6:**
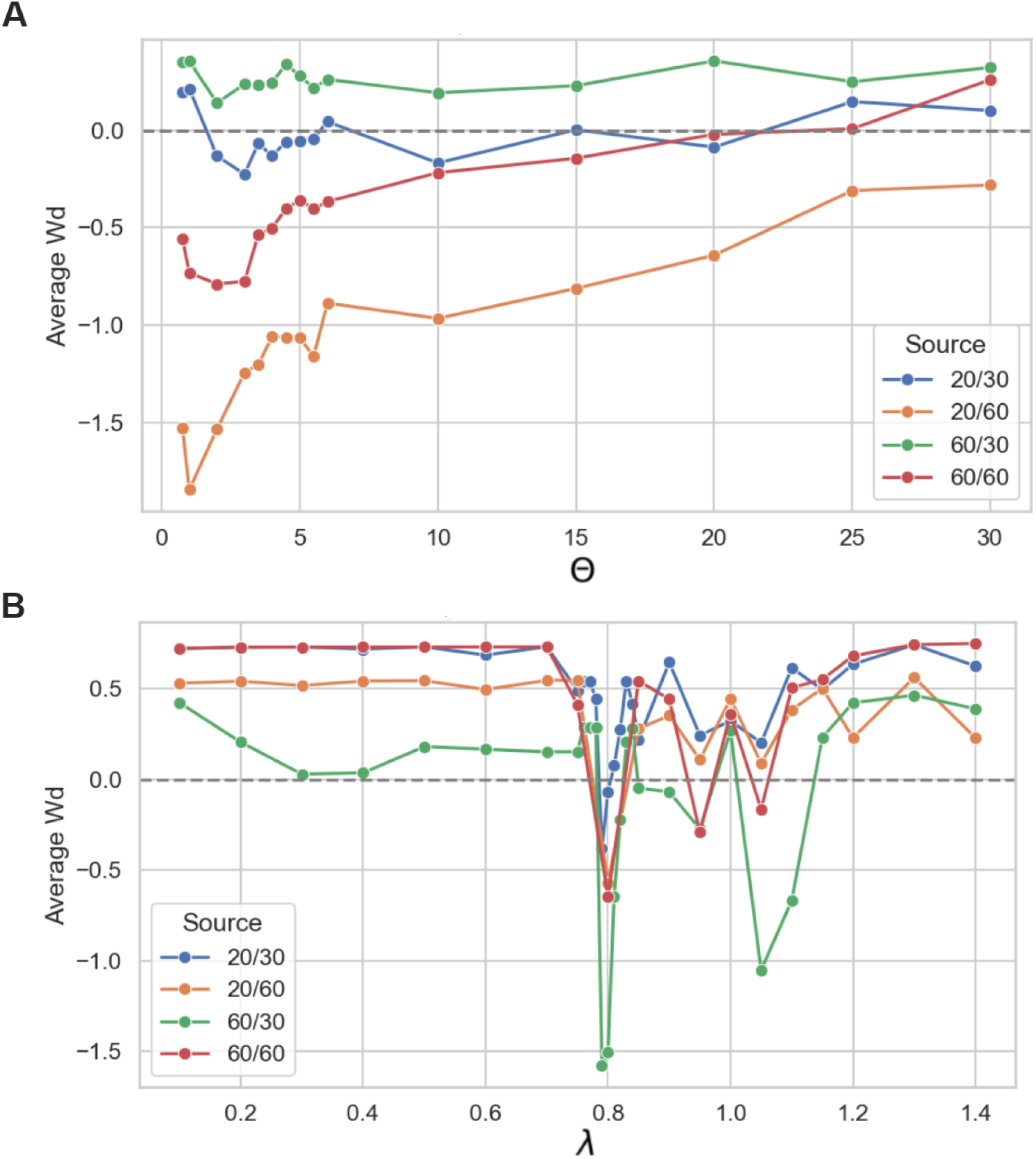
X-bit memory task performance across CA reservoir configurations. **(A)** *Sigmoid update.* Mean weighted score *W_d_* versus sigmoid gain Θ for four input–cue source geometries, denoted 20/30, 20/60, 60/30, and 60/60. The dashed horizontal line marks *W_d_* = 0. Performance varies systematically with source geometry and gain, with a shallow trough at intermediate Θ and recovery at larger Θ. **(B)** *Logistic–step update.* Mean weighted score *W_d_* versus relaxation rate *λ* for the same four source geometries. In contrast to the sigmoid case, all curves exhibit a pronounced minimum near the hard-threshold regime *λ* ≈ 0.8–1.0, followed by recovery away from this region. Curves show means over repeated trials with identical randomisation protocols; error bars are omitted for clarity. All runs used periodic boundaries and the same task protocol described in Sec. 3.1 and Methods Sec. 6.2.

Figure 7 refines this picture by showing how recall depends on the spatial interaction geometry (*R_a_, R_i_*). Across both update families, the strongest performance is concentrated in regions with broader inhibition than activation, typically *R_a_ < R_i_*, while excessive inhibition or overly local coupling reduces recall. For the sigmoid family (Figure 7A), the favourable region is comparatively smooth, indicating a relatively broad basin of useful operation across source geometries. For the logistic–step family (Figure 7B), the same qualitative preference for *R_a_ < R_i_* remains, but the landscape is more sharply structured and more sensitive to source geometry, consistent with the narrower operating window seen in the line-plot summary.

**Figure 7:**
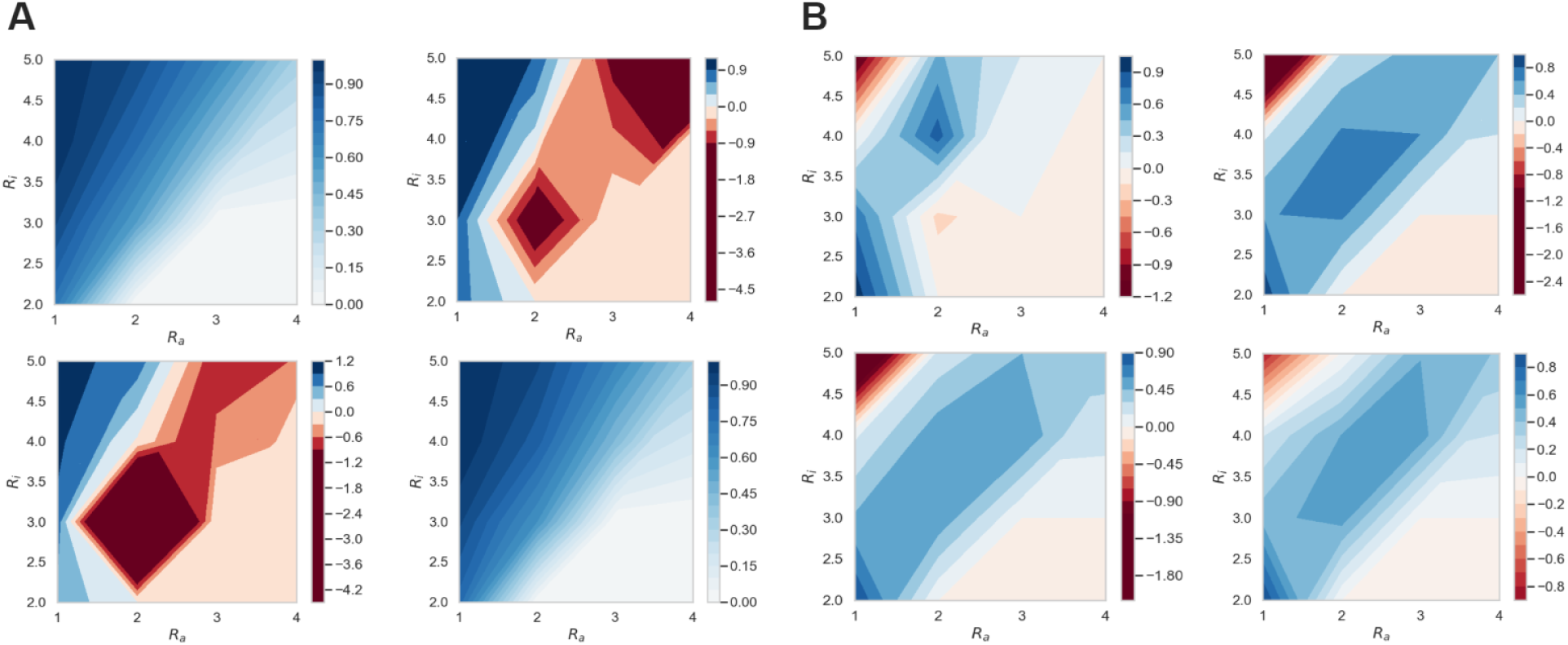
X-bit memory task: radius–radius performance landscapes across source geometries. **(A)** *Sigmoid update.* Mean weighted score *W_d_* over activator and inhibitor radii (*R_a_, R_i_*) for the four input–cue source geometries, using the same X-bit protocol as Figure 6. Across geometries, the strongest performance is concentrated in regions with broader inhibition than activation, typically *R_a_ < R_i_*, while overly local coupling or excessive inhibition reduces recall. **(B)** *Logistic–step update.* Corresponding radius–radius landscapes for the logistic–step family. The same broad preference for *R_a_ < R_i_* is retained, but the high-performing region is more sharply structured and more sensitive to source geometry, consistent with the narrower operating window seen in Figure 6B. Higher values indicate better delayed recall. All panels use the X-bit protocol of Sec. 3.1 and Methods Sec. 6.2, with identical randomisation settings across repeats.

Taken together, the X-bit results support three main conclusions. First, both update families exhibit a clearly structured dependence of recall on their tuning parameter rather than a flat response across settings. Second, spatial injection geometry modulates the attainable score but does not erase the family-specific performance profile. Third, the best-performing regimes are intermediate ones that preserve fading memory and transient separability without collapsing to trivial fixed points or rigid thresholded states. These findings are consistent with earlier CA-reservoir studies showing that useful temporal computation emerges in regimes balancing stability and separability rather than at either extreme [46–49]. Together with the image-classification results, they position the activator–inhibitor CA as a compact, tunable substrate for temporal computation.

### 3.2 Image classification

We evaluate the cellular-automaton reservoir as a fixed feature extractor for MNIST image classification. Each grayscale input image *u ∈* [0, 1]^28^*^×^*^28^ is quantised into eight binary bitplanes 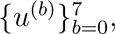 ordered from most-significant to least-significant bit. For each bit-plane, the current implementation embeds the 28 *×* 28 pattern into an *L_d_ ×L_d_* lattice with periodic boundaries by random spatial placement. When rep = 1, each bit-plane is embedded once into the full lattice; when rep *>* 1, the lattice is partitioned into a rep *×* rep grid of tiles and the same bit-plane is embedded at an independent random valid position inside each tile. No flips or rotations are used in the final CUDA pipeline.

Each embedded bit-plane then evolves for *T* iterations under the activator–inhibitor local balance *B*(*i, j, t*) from Eq. (3), using square activator and inhibitor neighbourhoods with radii (*R_a_, R_i_*), positive weights (*W_a_, W_i_*), and the logistic–step update rule from Eq. (8). At each iteration, the wrapped neighbourhood sums are computed by convolution on the torus, and the state is updated according to the sign of the local balance. The classification control parameter is the logistic–step relaxation rate *λ_b_*, which governs contraction for 0 *< λ_b_ <* 1, the hard-threshold limit at *λ_b_* = 1, and overshoot for *λ_b_ >* 1.

Unless noted otherwise, 2 *×* 2 max-pooling is applied after each iteration. Flattened pooled snapshots from all eight bit-planes and all *T* time steps are concatenated to form the final feature vector

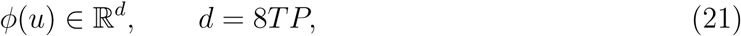

where *P* = (*L_d_*/*s*)^2^ when pooling with factor *s* is used and 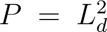 otherwise. Thus, the CA is treated here as a fixed nonlinear map from image space to a high-dimensional spatio-temporal feature representation; only the readout is trained. The purpose of the MNIST experiment is therefore not to optimise image-classification performance, but to test whether the nonlinear medium transforms structured inputs into representations that are linearly decodable across dynamical regimes.

Classification is performed by first standardising the feature matrix with `StandardScaler` and then fitting a multinomial logistic-regression readout with L-BFGS. We therefore interpret the MNIST experiment as an empirical test of how well the untrained activator–inhibitor dynamics transform digit images into linearly separable features, rather than as a setting in which the CA itself is optimised by gradient descent.

Figure 8 summarises performance for four sample sizes, *N ∈ {*200, 500, 1000, 2000*}*. For each run, a fresh subset of size N is sampled uniformly without replacement, split into train/test partitions (80/20), decomposed into bit-planes, mapped through the CA reservoir, and classified by the logistic-regression readout. The blue curve reports the *mean aggregation*, i.e. the average accuracy at each *λ_b_* across the full sweep of (*R_a_, R_i_, W_a_, W_i_*). The red curve reports the *best-mean aggregation*, i.e. the best-performing setting at each *λ_b_* within each run, averaged across runs.

**Figure 8:**
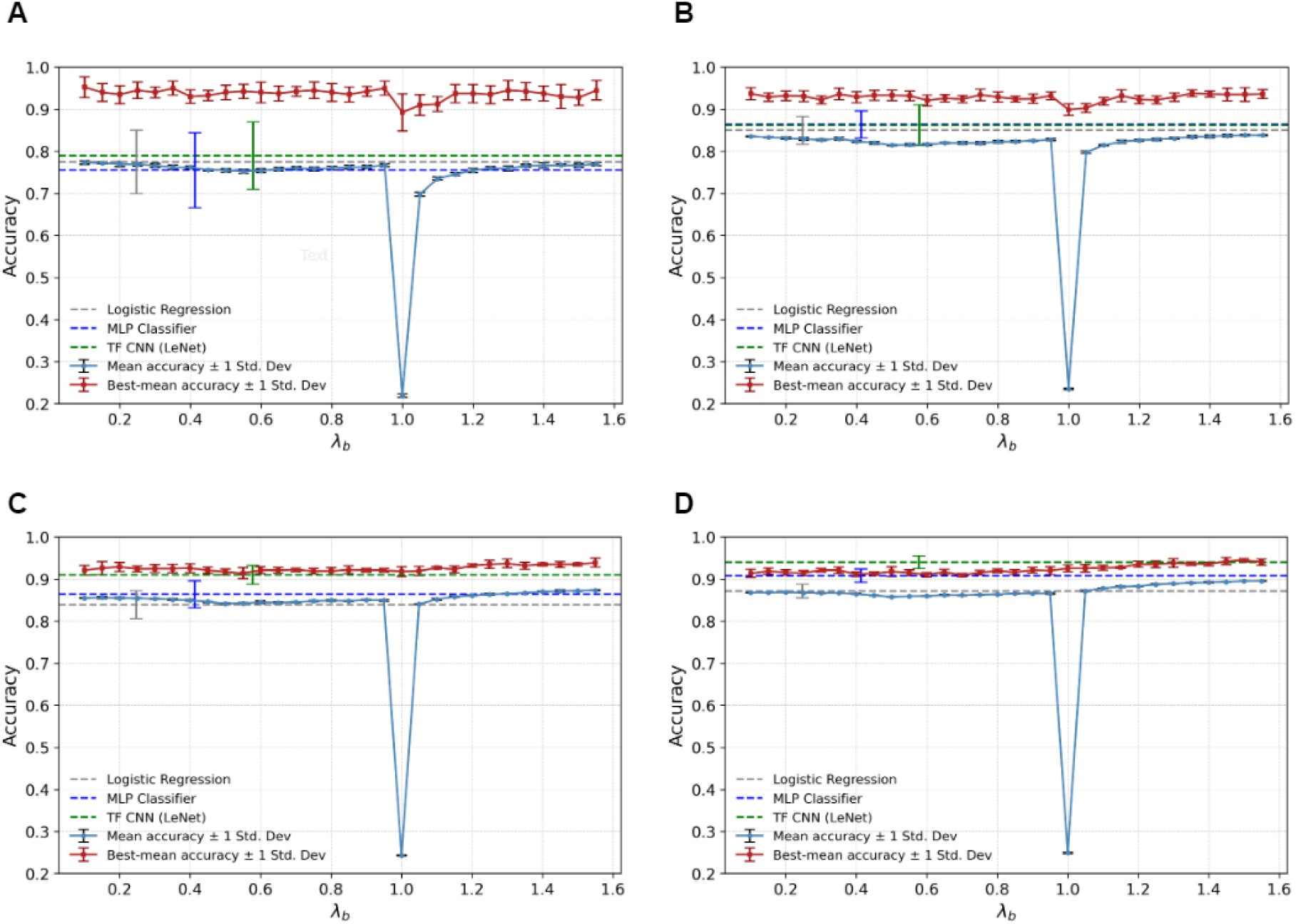
MNIST classification accuracy of the CA reservoir across sample sizes. Panels **(A)**–**(D)** show results for MNIST subsets of size *N* = 200, 500, 1000, and 2000, respectively. For each run, a fresh subset of size *N* is sampled, split into training and test sets (80*/*20, stratified), decomposed into eight bit-planes, evolved under the activator–inhibitor CA reservoir, and classified by a standardised multinomial logistic-regression readout. The blue curve shows the *mean aggregation*, i.e. the average accuracy at each *λ_b_* over the full sweep of (*R_a_, R_i_, W_a_, W_i_*). The red curve shows the *best-mean aggregation*, defined as the best-performing CA setting within each run at a given *λ_b_*, averaged over runs. Error bars denote ±1 standard deviation over 5 runs. Horizontal dashed lines indicate the corresponding benchmark accuracies for logistic regression, MLP, and TF CNN (LeNet) at the same sample size. Across all four panels, the mean curve remains comparatively smooth and improves with sample size, whereas the best-mean envelope stays consistently high, indicating strong hyperparameter sensitivity but reproducibly strong tuned regimes. A sharp collapse persists near *λ_b_*≈ 1, consistent with the hard-threshold limit of the logistic–step update.

Several features are robust across all four sample sizes. First, away from *λ_b_ ≈* 1, the mean curve is relatively smooth and improves gradually with increasing *N*, indicating that the typical quality of the CA-induced feature family increases with data scale. Second, the best-mean curve remains substantially above the mean curve, showing that strong performance is concentrated in a restricted but reproducible subset of operating points rather than being uniform across parameter space. Third, all four panels exhibit a pronounced collapse of the mean curve near *λ_b_ ≈* 1, indicating that the hard-threshold point is structurally pathological on average: most operating points fail there, while only a narrow viable subset remains.

This distinction between the two summaries is important. The mean curve is the more conservative descriptor of *typical* CA-reservoir behaviour under the chosen sweep, whereas the best-mean curve quantifies the attainable tuned envelope within the same run-specific split. Accordingly, the red curve should be interpreted as evidence that the CA reservoir contains high-performing regimes, not as evidence that the entire reservoir family uniformly reaches that performance level.

A complementary view is provided by the contour analysis in Fig. 9, which plots the best-mean accuracy surface over (*R_a_, R_i_*) at the empirically selected *λ^∗^* for each sample size. These surfaces show that the strongest MNIST regimes are not diffuse across parameter space, but cluster in a compact band of moderate radii, typically with *R_i_* ≳ *R_a_*. Thus, the MNIST results support the same broader conclusion as the tuning and memory experiments: the computational utility of the medium is strongly regime-dependent, and the best image-classification performance is obtained only in a restricted part of the activator–inhibitor parameter space.

**Figure 9:**
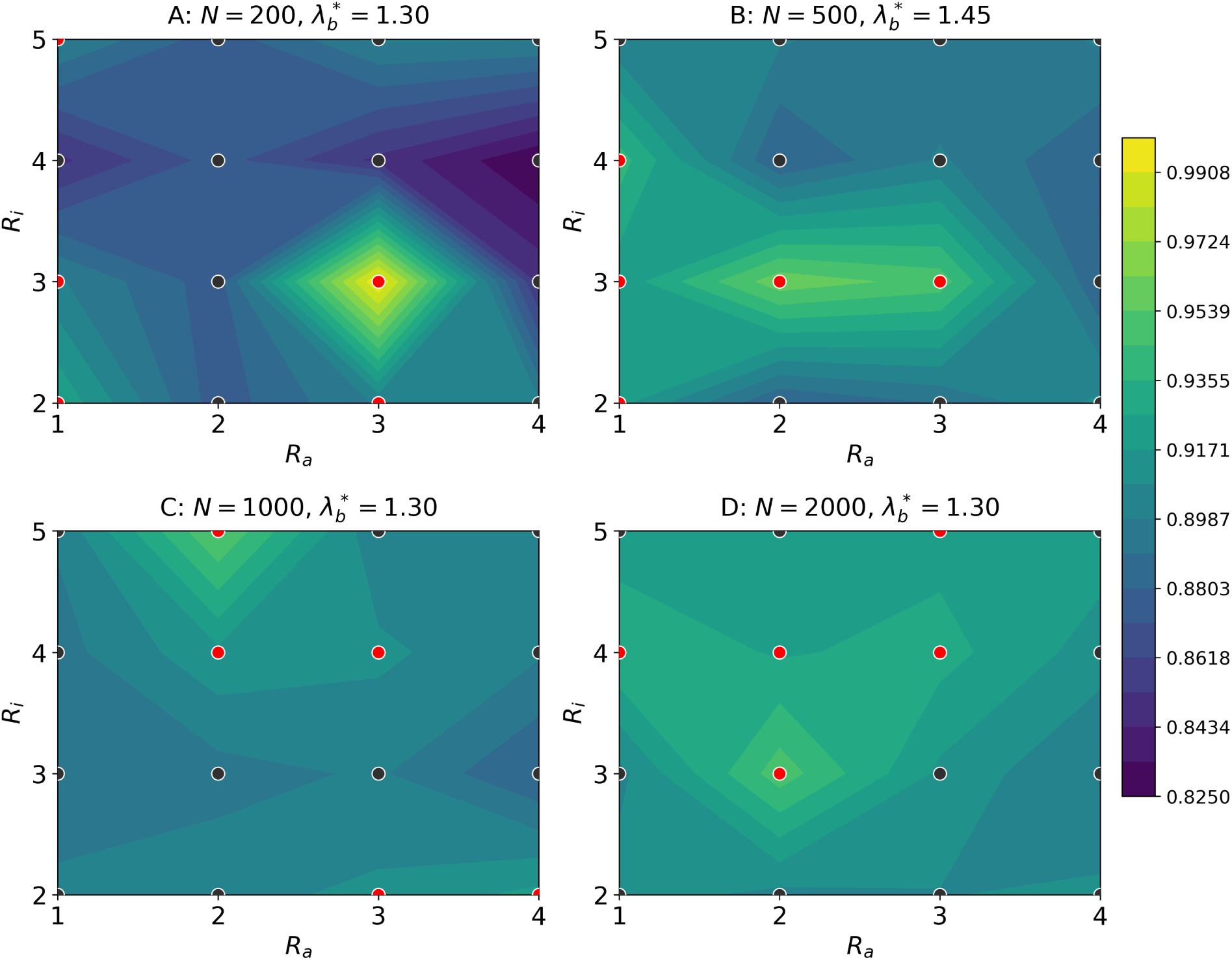
MNIST contour analysis of the CA reservoir across sample sizes. Panels **(A)**–**(D)** show the best-mean test-accuracy surface over activator and inhibitor radii (*R_a_, R_i_*) for MNIST subsets of size *N* = 200, 500, 1000, and 2000, respectively. For each sample size, the update rate *λb*^∗^ is selected as the value maximising the run-averaged best accuracy across the full sweep of (*R_a_, R_i_, W_a_, W_i_*); the selected values are reported in the panel titles. At this fixed *λ*_b_*, each surface is constructed by taking, for each run and each (*R_a_, R_i_*) pair, the best accuracy over (*W_a_, W_i_*) and then averaging these best values across runs. Across sample sizes, the strongest-performing regimes cluster in a compact band of moderate radii, typically with *R_i_* ≳ *R_a_*, rather than being diffuse across parameter space. This supports the conclusion that MNIST performance is strongly regime-dependent and is concentrated in a restricted geometric region of the activator–inhibitor parameter space.

## 4 Discussion

This work provides a nonlinear-regime interpretation of computation in an activator–inhibitor cellular automaton. A single local interaction rule, combined with tunable activation or relaxation, generates distinct dynamical behaviours in which pattern complexity, fading memory, and input separability vary together, but not in a simple one-to-one manner. Reservoir computing is used here as a probe of these behaviours, revealing when self-organised spatiotemporal transients remain rich enough to retain information and structured enough to support a linear readout.

The value of the system is not that it outperforms specialised digital machine-learning architectures. Rather, it shows how a nonlinear spatial medium can be moved between different computational regimes while keeping the local rule interpretable. The same activator–inhibitor wiring supports collapsed, structured, saturated, and overshooting dynamics, and these behaviours differ in their ability to retain perturbation history and separate inputs for downstream readout. The model therefore sits naturally alongside physical reservoir-computing approaches, with the useful feature that the relevant control parameters, including gain, relaxation, interaction radii, and coupling weights, remain explicit.

Taken together, the results indicate that computation in this medium is regime-dependent, rather than a generic property of the update rule. The tuning curves, radius-by-radius landscapes, and task results show that useful computation is concentrated in restricted operating regions instead of being spread uniformly across parameter space. In both the X-bit and MNIST experiments, the strongest regimes occur away from the hard-threshold limit and typically favour broader inhibition than activation, with moderate radii producing the most useful spatiotemporal transients. In this system, the “edge of chaos” idea is therefore best understood not as a preference for maximal irregularity, but as a structured dynamical regime in which fading memory and linear separability are both preserved.

DNOS and DPC are best interpreted as operational diagnostics of this regime structure. They do not establish a universal relationship between complexity and performance, but they do capture broad transitions that also appear in the task results. Collapsed or frozen states tend to reduce both diagnostic richness and computational utility, whereas regimes that preserve multilevel transients and mobile fronts are more often associated with improved recall and classification. Reservoir computing therefore serves here primarily as a probe of morphogenetic state space, identifying when the untrained medium generates internal representations that remain useful for downstream readout.

Several limitations should be clear. First, the present study provides a computational regime map rather than a hardware implementation, so claims about energy efficiency, latency, or device-level advantage remain prospective. Second, the MNIST task is used as a diagnostic of separability, not as evidence of leading image-classification performance. Third, the cellular automaton is inspired by reaction–diffusion dynamics, but is not analysed here through formal Turing instability criteria. The relevant claim is therefore about nonlinear regime structure under local activation and inhibition, not strict PDE-based Turing pattern classification. Finally, DNOS and DPC are operational coarse-grained diagnostics whose numerical values depend on the stated discretisation and compression protocol, although their role here is comparative rather than absolute.

The results also point to a cautious biological and physical-computing hypothesis in which effective gains, thresholds, and relaxation rates may act not only as patterning controls, but also as controls on computational regime. From this perspective, tuning a tissue-like or physical nonlinear medium could balance rapid stabilisation against the retention of perturbation history. Future work could test this idea more directly using adaptive tuning rules, hardware implementations, and complementary separability measures, such as intrinsic separation dimension, to connect dynamical regime structure more explicitly to learnability.

## 5 Methods

### 5.1 CA pattern-formation parameter sweeps with sigmoid and logistic–step activations

All simulations were implemented in Python 3.10 and C++17. The workflow here underpins the results in Sec. 2.4. Model equations are defined in the main text (Eqs. 3–8; Sec. 2.1). The computational pipeline follows Algorithm 1, with initialisation, stepping, and local-balance evaluation handled by Algorithms 2, 3, and 4, respectively.

#### Sweep design

We systematically explored activator/inhibitor radii, couplings, and a single family-specific tuning parameter. Radii were drawn from *R_a_* ∈ {1, 2, 3, 4} and *R_i_* ∈ {1, 2, 3, 4, 5}; weights spanned *W_a_, W_i_* ∈ {0.1, 0.2, . . ., 1.0}. For the *sigmoid* family (Eq. 5), the gain Θ was swept over [0.1, 30.0]; for the *logistic–step* family (Eq. 8), the relaxation rate *λ* was swept over [−3.0, 5.35] with finer sampling near 0 and 1. Each configuration (*R_a_, R_i_, W_a_, W_i_,* Θ) or (*R_a_, R_i_, W_a_, W_i_, λ*) was repeated for *N*_rep_=10 random seeds (Algorithm 1).

#### Simulation protocol

Runs used a square lattice *L* of size *S×S* with periodic boundaries (*S*=80 by default). At *t*=0, cells were initialised by assigning *x_i,j_*(0) *∼ U*(0, 1) with probability *p*_seed_=0.5, and 0 otherwise (Algorithm 2). Synchronous updates employed a double buffer (Algorithm 3). At each iteration, activator and inhibitor contributions were computed from nested Moore neighbourhoods of radii (*R_a_, R_i_*), and states were updated via either the sigmoid rule or the logistic–step rule using the local balance (Eqs. 3, 5, 8; Algorithms 3, 4). All arithmetic used `float32` for hardware consistency. Typical trajectory lengths were T =60 (sigmoid) and T =30 (logistic–step).

#### Complexity measurement

After the final iteration, DNOS and DPC were computed using the current notebook implementation rather than a single shared fixed-bin discretisation. For DNOS, each final lattice was first checked for values outside [0, 1]; if necessary, `NormalizeData` was applied. The lattice was then mapped to the precision-controlled array around(10*^p^X, p*), and DNOS was recorded as the number of unique entries (Eq. 11; Algorithm 5). For DPC, compression ratios were computed for all generated images, assembled into a matrix with one row per parameter-setting ensemble, globally normalised over all entries with `NormalizeData`, digitised into *n_b_* equally spaced bins on [0, 1], and reduced row-wise to the number of occupied bins (Eq. 14; Algorithm 6). Intermediate artefacts—final state matrices, compression-ratio tables, and DNOS/DPC summaries—were saved as CSV files for reproducibility.

#### Execution and logging

C++ kernels were parallelised with OpenMP; Python orchestration used `joblib`. Each run produced a JSON metadata record (parameters, seed, iteration count, DNOS, DPC). Aggregated tables were used to generate tuning curves (Figure 2), radius–radius landscapes (Figure 3), and state-space scatter plots with exemplar histograms (Figs. 4–5). All experiments used the environment in Sec. 5.4.

### 5.2 Reservoir computing task: X-bit memory

We assessed the information–storage capacity of the CA reservoir using the standard X-bit recall benchmark [46, 47]. The objective is to reproduce the input bit presented *X* steps earlier, thereby quantifying fading memory. The overall protocol is summarised in Algorithm 7; data injection and readout training are detailed in Algorithms 8 and 9.

#### Task and data injection

For each trial, a binary input stream was presented over four channels (signal, flipped signal, distractor, cue). At every step, inputs were injected by overwriting a randomly permuted subset of lattice sites on an *L_d_ × L_d_* torus (periodic boundaries) using the randomised write procedure in Algorithm 8. Unless stated otherwise, a write probability `prob∈` (0, 1) controlled the fraction of updated sites per step, and a short binary token of length `STRZ`= 4 was written to produce overlapping yet non-identical activation patterns across time. After injection, the lattice evolved for *I* reservoir updates under the activator–inhibitor balance

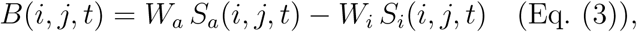

with square neighbourhood radii (R*_a_*, ℝ*_i_*) and couplings (W*_a_*, W*_i_*) > 0 (short–range activation, longer–range inhibition), implemented via the generic stepping routine (Algorithm 3) calling the local-balance kernel (Algorithm 4).

#### Update families (fixed wiring)

Both continuous–state updates defined earlier were tested on the same *B*(*i, j, t*): (i) the *sigmoid rule* (Eq. 5), where the gain Θ controls instantaneous nonlinearity, and (ii) the *logistic–step relaxer* (Eq. 8), where the rate *λ* sets relaxation/overshoot. Thus, tuning acts on nonlinearity (Θ) or memory timescale (*λ*) without altering local wiring (Algorithm 3).

#### Readout and metric

At each iteration *t*, the flattened lattice state **x***_t_* formed the reservoir feature vector recorded by Algorithm 7. A linear readout *W*_out_ (ridge regression; Algorithm 9) was trained to reconstruct the delayed input upon cue presentation. Performance was reported by the weighted X-bit score *W_d_* in Eq. 20, averaged over trials and delays, following [46, 47].

#### Sweep settings

We scanned (*R_a_, R_i_*) *∈ {*2*, . . .,* 5*}*^2^ and (*W_a_, W_i_*) on a coarse grid subject to *R_a_ < R_i_* and *W_a_ > W_i_*. For the logistic–step family we densely swept *λ* over [0.2, 1.4] (main results); for the sigmoid family we varied the gain Θ over the operating range used in Sec. 2.4. Four input–cue geometries (“20/30”, “60/30”, “20/60”, “60/60”) probed sensitivity to spatial write patterns. All runs used identical randomisation protocols across repeats. Extended sweeps exploring *λ <* 0 and *λ >* 1 (overshoot) and additional lattice sizes *S ∈* {30, 50, 80} are reported in the Supplement. Sweep orchestration followed Algorithm 1.

#### Implementation notes and complexity linkage

Simulators produced per–step state trajectories that were standardised and fed to the readout (Algorithms 7, 9). DNOS and DPC were computed per configuration (Algorithms 5–6) using the same coarse–graining as in Sec. 5.1, enabling correlation analyses between complexity measures and memory scores *prior* to readout training.

##### Reproducibility

Code compiled from the C++ cores for the two update families (sigmoid / logistic–step) executed synchronous updates over (*R_a_, R_i_*) neighbourhoods (Algorithm 3). Each configuration was run for ten seeds in the environment described in Sec. 5.4; outputs and trained readouts are archived under `cpp_results/`.

### 5.3 Reservoir computing for image classification (MNIST)

We assess the CA reservoir on MNIST using the current CUDA implementation of the bit-plane reservoir pipeline and the logistic–step dynamics defined in Eq. (8). The feature-extraction and sweep protocol are given in Algorithms 10–12.

#### Experimental setup

Each grayscale image *u ∈* [0, 1]^28^*^×^*^28^ is quantised into eight Boolean bit-planes 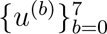 (most-significant to least-significant bit). For each bit-plane, the current implementation embeds the 28 *×* 28 pattern into an *L_d_ × L_d_* lattice with toroidal boundaries by random spatial placement. When `rep` = 1, each bit-plane is embedded once into the full lattice; when `rep >` 1, the lattice is partitioned into a `rep × rep` grid of tiles and the same bit-plane is embedded at an independent random valid position inside each tile. No flips or rotations are used in the final CUDA pipeline.

For each embedded bit-plane, the lattice evolves for *T* iterations under square activator and inhibitor neighbourhoods of radii (*R_a_, R_i_*) and positive weights (*W_a_, W_i_*). At each step, the local balance term is computed from wrapped convolutional neighbourhood sums, and the state is updated by the logistic–step rule

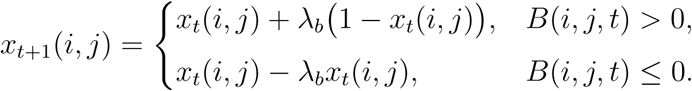

Unless noted otherwise, 2*×*2 max-pooling is applied after each iteration. Flattened snapshots from all eight bit-planes and all *T* time steps are concatenated to form the final feature vector

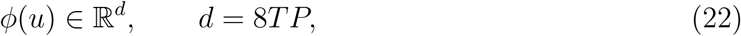

where P = (L*_d_*/s)^2^ when pooling with factor s is enabled and 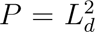 otherwise. Features are standardised with `StandardScaler`, and classification is performed with a multinomial logistic-regression readout trained with L-BFGS (`max_iter`=2000).

#### Evaluation protocol

For each sample size *N ∈ {*200, 500, 1000, 2000*}* and each of *R* = 5 runs, a subset of *N* MNIST images is sampled uniformly without replacement and split into train/test partitions using a stratified 80:20 split. Bit-plane decompositions are then precomputed once for the training and test subsets of that run and reused across the full hyperparameter sweep.

The CA sweep spans

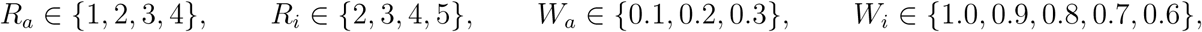

together with a scan over update rates *λ_b_* from 0.10 to 1.55 in steps of 0.05. For each (*R_a_, R_i_, W_a_, W_i_, λ_b_*) combination, CA features are extracted on the training and test sets, standardised, and evaluated with multinomial logistic regression. Test accuracy is logged for every run and every grid point.

Figure 8 reports two summaries at each *λ_b_*: (i) *mean aggregation*, defined as the mean accuracy across all swept CA settings within a run, followed by mean *±* s.d. across runs; and (ii) *best-mean aggregation*, defined as the best-performing CA setting within a run at that *λ_b_*, followed by mean *±* s.d. across runs. Panels **(A)**–**(D)** correspond to sample sizes *N* = 200, 500, 1000, and 2000, respectively.

Figure 9 provides a complementary contour analysis over (*R_a_, R_i_*). For each sample size, the optimal update rate *λ^∗^* is first selected as the value maximising the run-averaged best accuracy across the full sweep of (*R_a_, R_i_, W_a_, W_i_*). At this fixed *λ^∗^*, the contour surface is obtained by taking, for each run and each (*R_a_, R_i_*) pair, the best accuracy over (*W_a_, W_i_*) and then averaging these best values across runs.

#### Baselines

For each sample size, horizontal reference bands in Figure 8 show mean *±* s.d. accuracies over the same number of runs for three non-reservoir baselines: (a) logistic regression on raw pixels, (b) an MLP classifier with hidden sizes (1024, 512, 128), and (c) a compact LeNet-style CNN consisting of two convolution/max-pooling blocks (32 filters of size 3 *×* 3, then 64 filters of size 3 *×* 3), followed by a dense layer of width 64 and a softmax output layer. The CNN baseline is trained with Adam for 20 epochs using batch size 128; all baselines are evaluated under the same sample-size and repeated-run protocol as the CA reservoir.

### 5.4 Computational environment and reproducibility

#### Hardware and execution

The parameter-sweep and X-bit experiments were executed with the C++/Python pipeline described above, whereas the MNIST reservoir experiments used a GPU-enabled PyTorch implementation that performs batchwise CA evolution and explicitly frees cached GPU memory during the sweep. The MNIST implementation also constrains CPU thread oversubscription by setting OpenMP, MKL, OpenBLAS, and NumExpr thread counts to one. Baseline CNN experiments were run in TensorFlow with memory-growth enabled when a GPU was available; otherwise, they fall back to CPU.

#### Determinism and randomisation

The current CUDA MNIST sweep uses a fixed base seed and derives per-run random number streams from that seed. Consequently, image sampling, stratified train/test splitting, and the random embedding locations are reproducible for a given run index and sweep order. Within each run, the same sampled subset and train/test split are reused across the full (*R_a_, R_i_, W_a_, W_i_, λ_b_*) sweep. By contrast, the baseline scripts may use either fixed or fresh seeds depending on configuration; unless otherwise stated, reported baseline statistics are means and standard deviations over 5 independently repeated runs at each sample size.

#### Reproducibility for MNIST aggregation

All MNIST CA runs log per-configuration test accuracies to CSV files containing run, `lb, Ra, Ri, Wa, Wi`, and accuracy. For the blue curve in Figure 8, accuracies are grouped by λ*_b_*, averaged across CA settings within each run, and then averaged across runs (mean ± s.d.). For the red curve, the best CA setting at each *λ_b_* is selected within each run before averaging across runs (best-mean ± s.d.). This makes the blue curve a descriptor of typical performance across the sweep, whereas the red curve is a tuned upper envelope for the same run-specific split. For Figure 9, *λ^∗^* is selected separately for each sample size from the best-mean curve, and the (*R_a_, R_i_*) surface is then constructed by taking the best accuracy over (*W_a_, W_i_*) within each run and averaging across runs.

#### Implementation details

The end-to-end MNIST workflow corresponds to Algorithms 10–12. The reservoir code uses wrapped convolutional neighbourhood sums, optional per-iteration max-pooling, and batchwise GPU feature extraction, while the readout is a convex multinomial logistic-regression classifier trained after feature standardisation.

#### Archival and reproducibility

The full computational framework, including the CUDA CA reservoir implementation, baseline scripts, parameter grids, CSV outputs, and figure-generation notebooks, is archived in the project repository. This allows deterministic reproduction of the seeded MNIST sweeps and statistical replication of the reported aggregated results.

##### Algorithm 1

Parameter sweep for CA with sigmoid or logistic–step updates

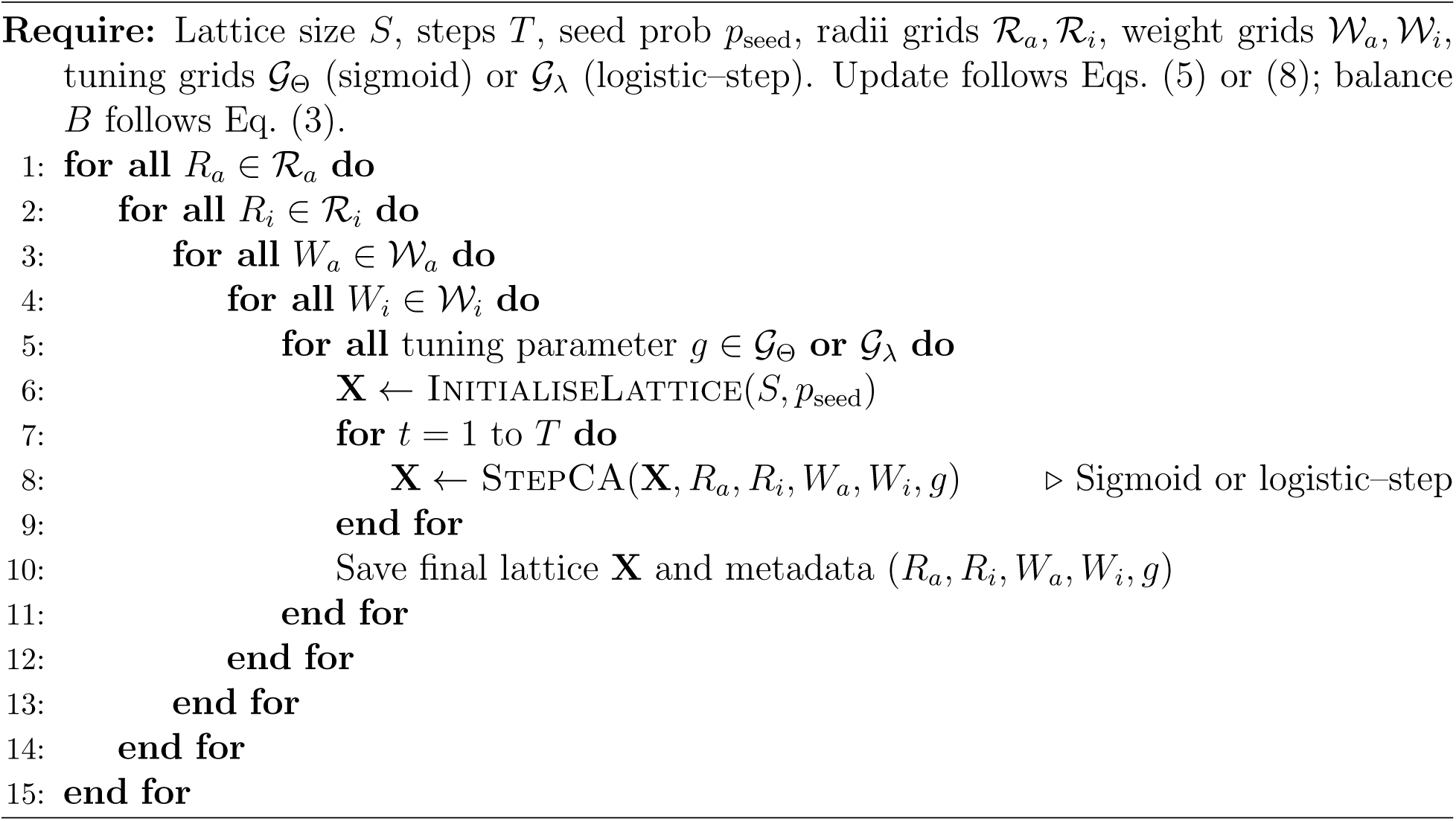

##### Algorithm 2

InitialiseLattice

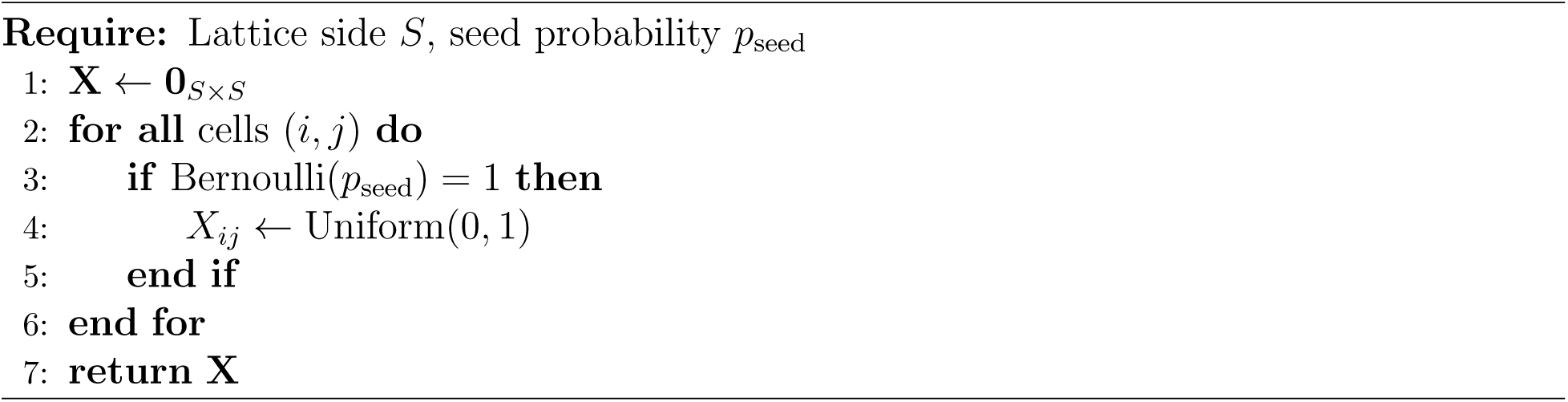

##### Algorithm 3

StepCA (generic wrapper)

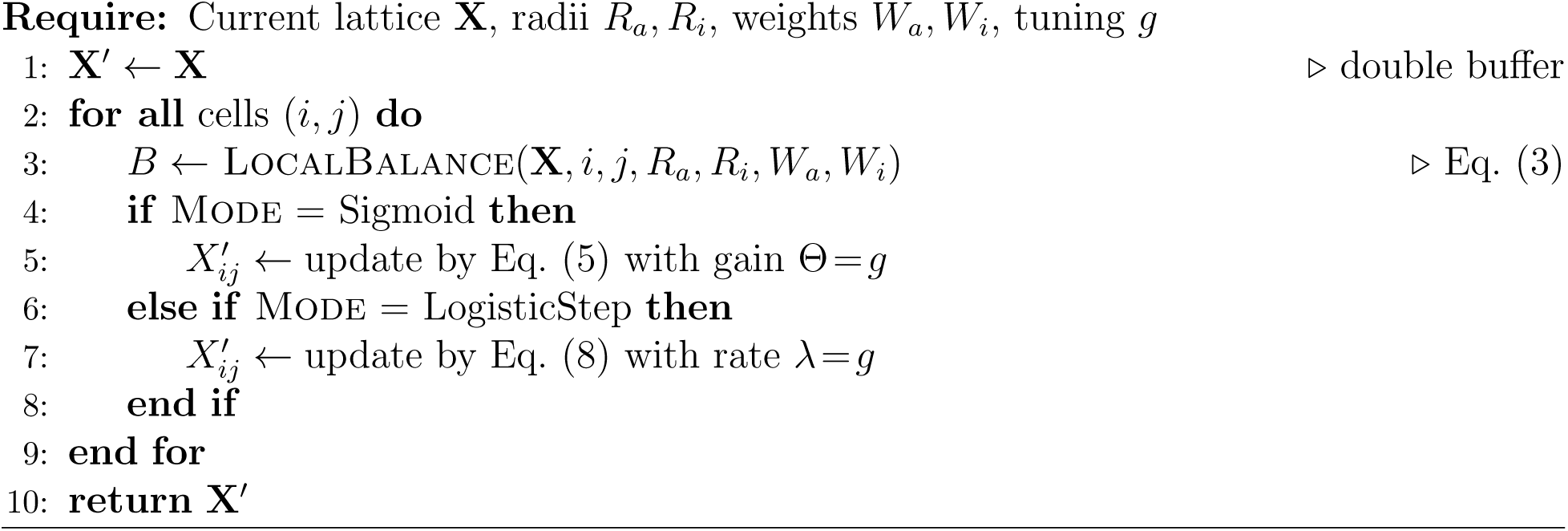

##### Algorithm 4

LocalBalance

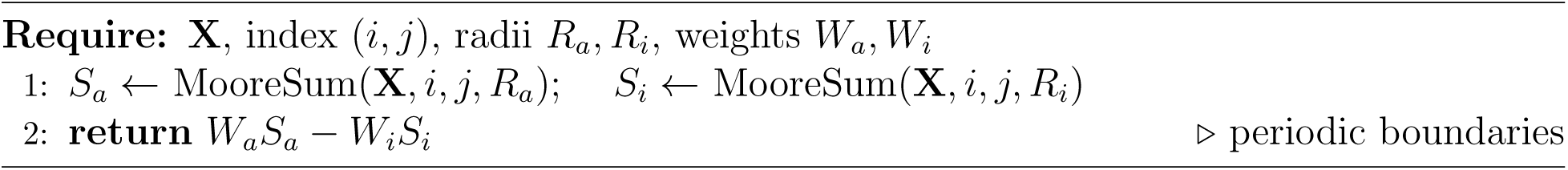

##### Algorithm 5

ComputeDNOS (alphabet breadth; current implementation)

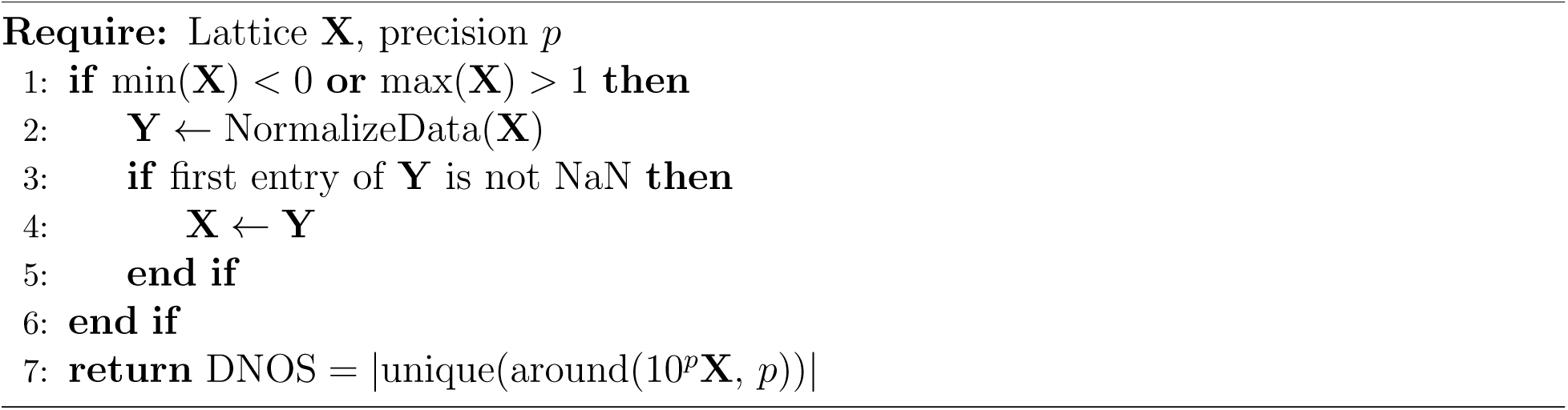

##### Algorithm 6

ComputeDPC (algorithmic depth across ensembles; current implementation)

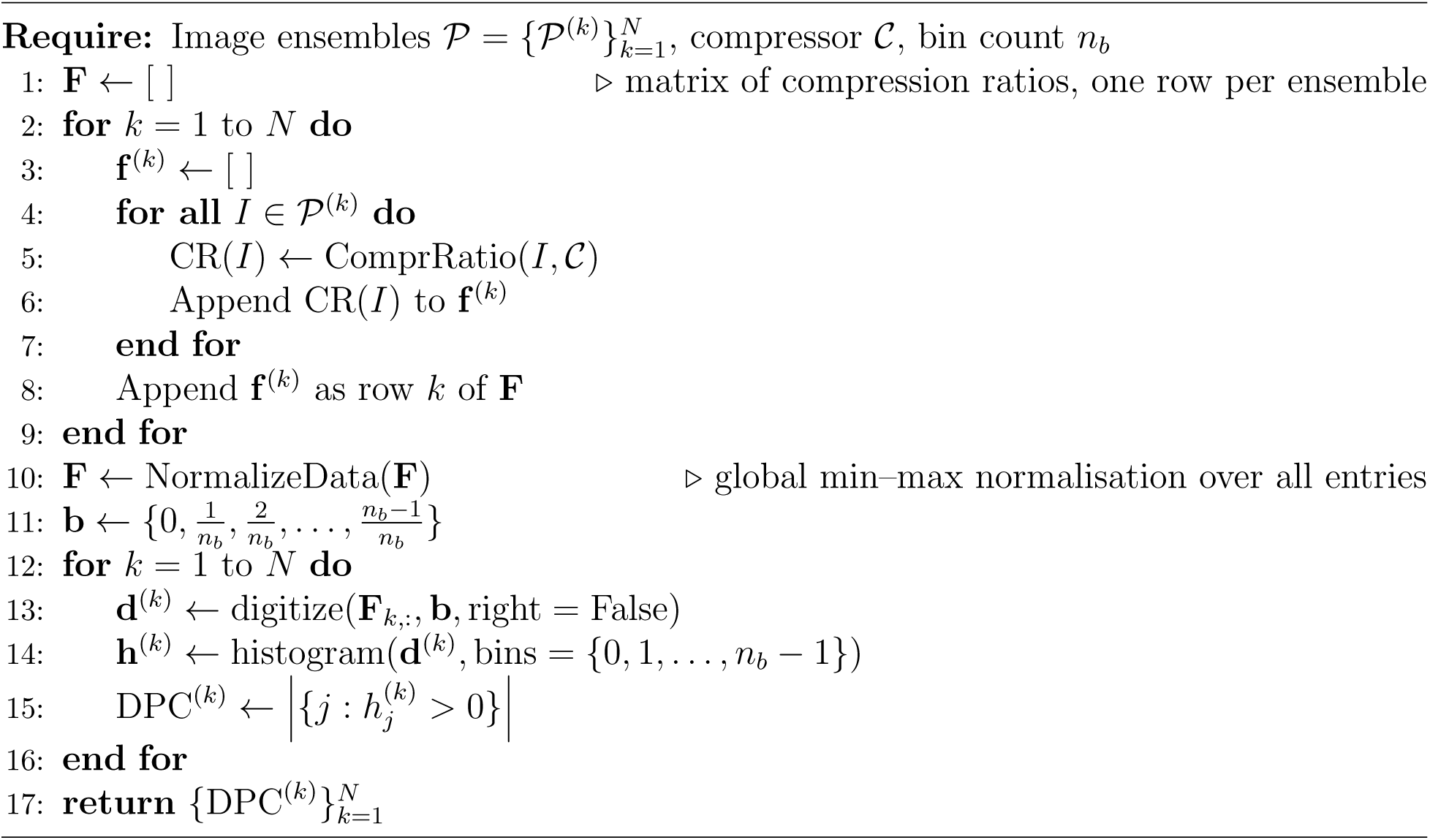

##### Algorithm 7

XBitMemoryTask

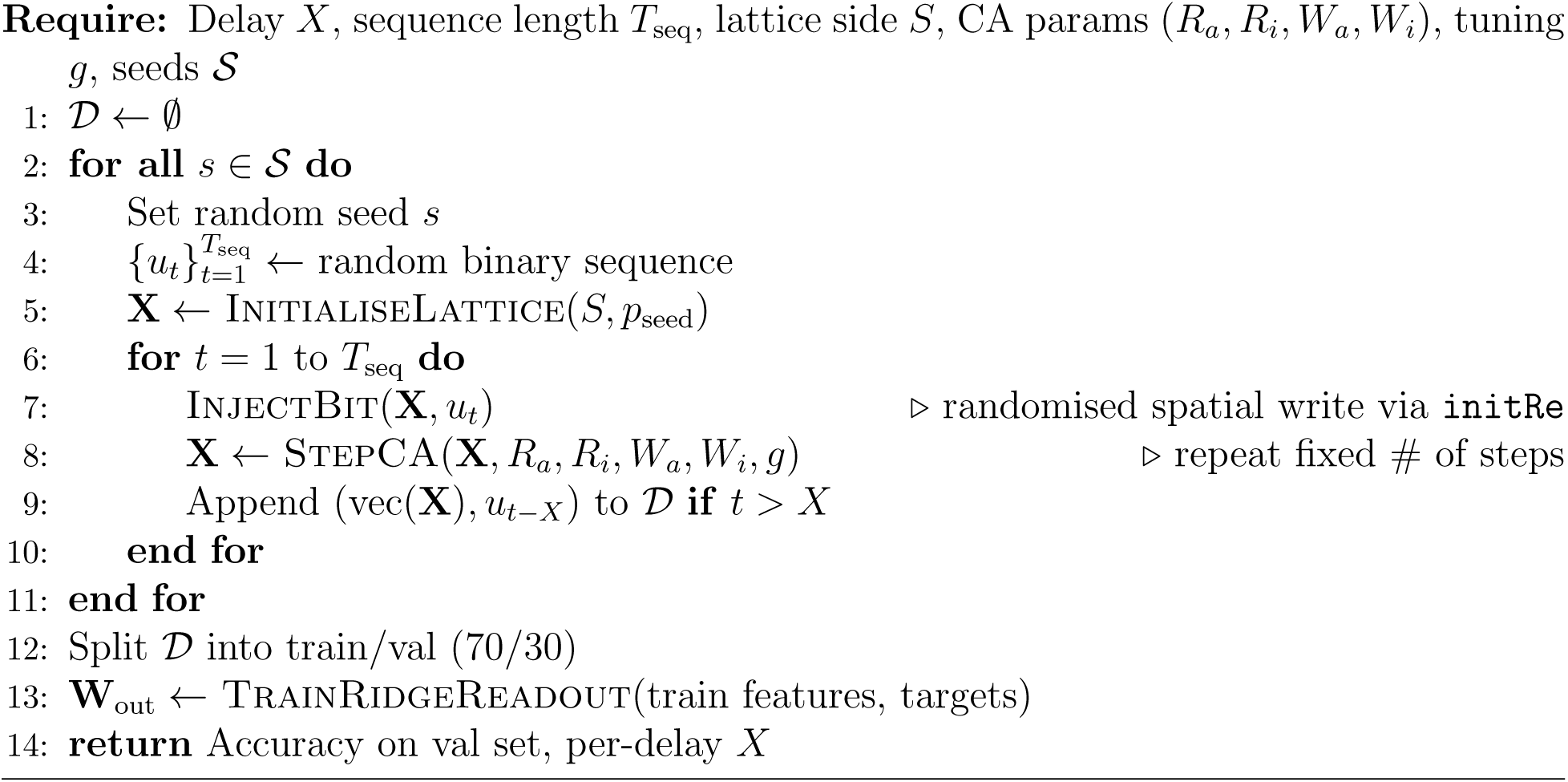

##### Algorithm 8

InjectBit

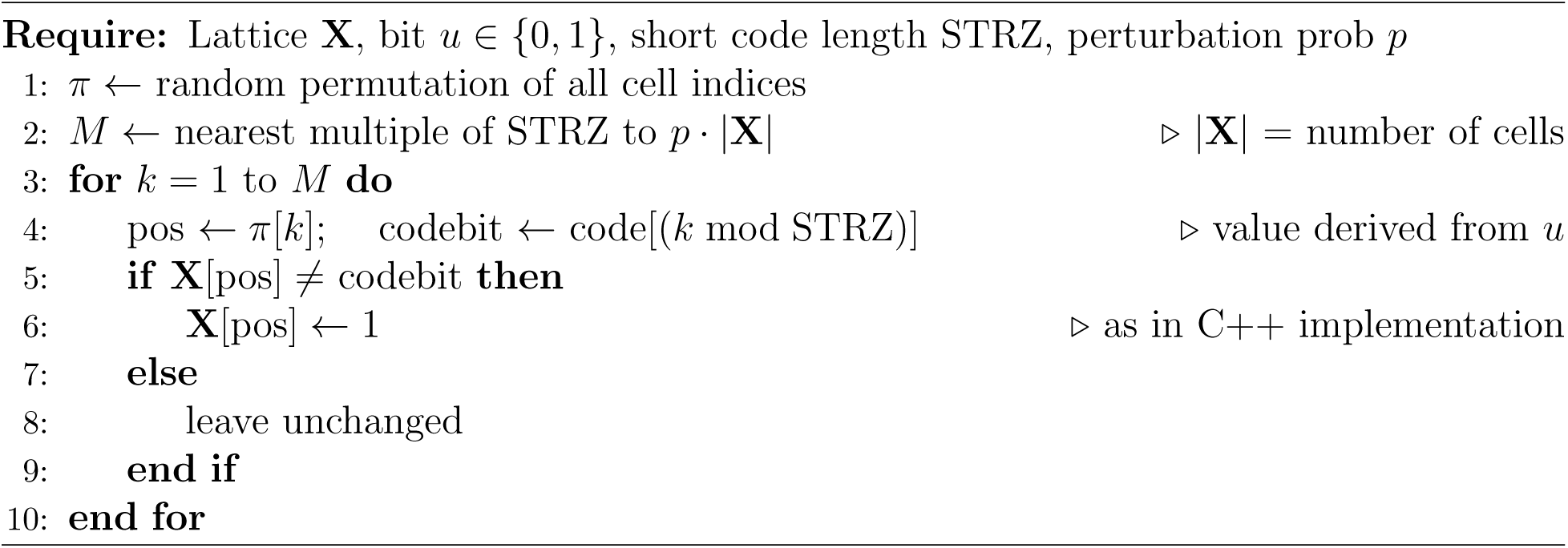

##### Algorithm 9

TrainRidgeReadout

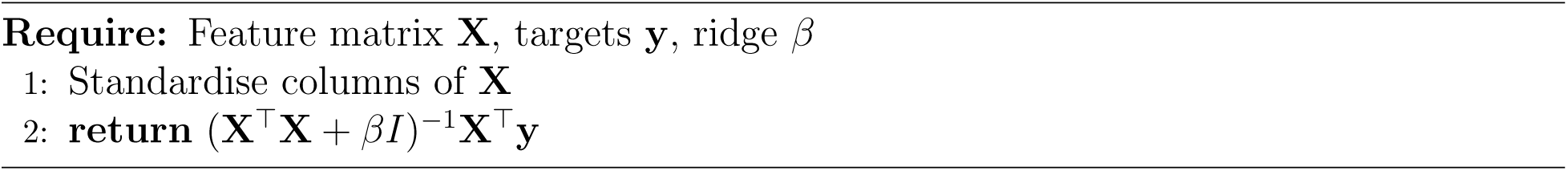

##### Algorithm 10

MNISTFeatureExtractionCUDA (current CA reservoir)

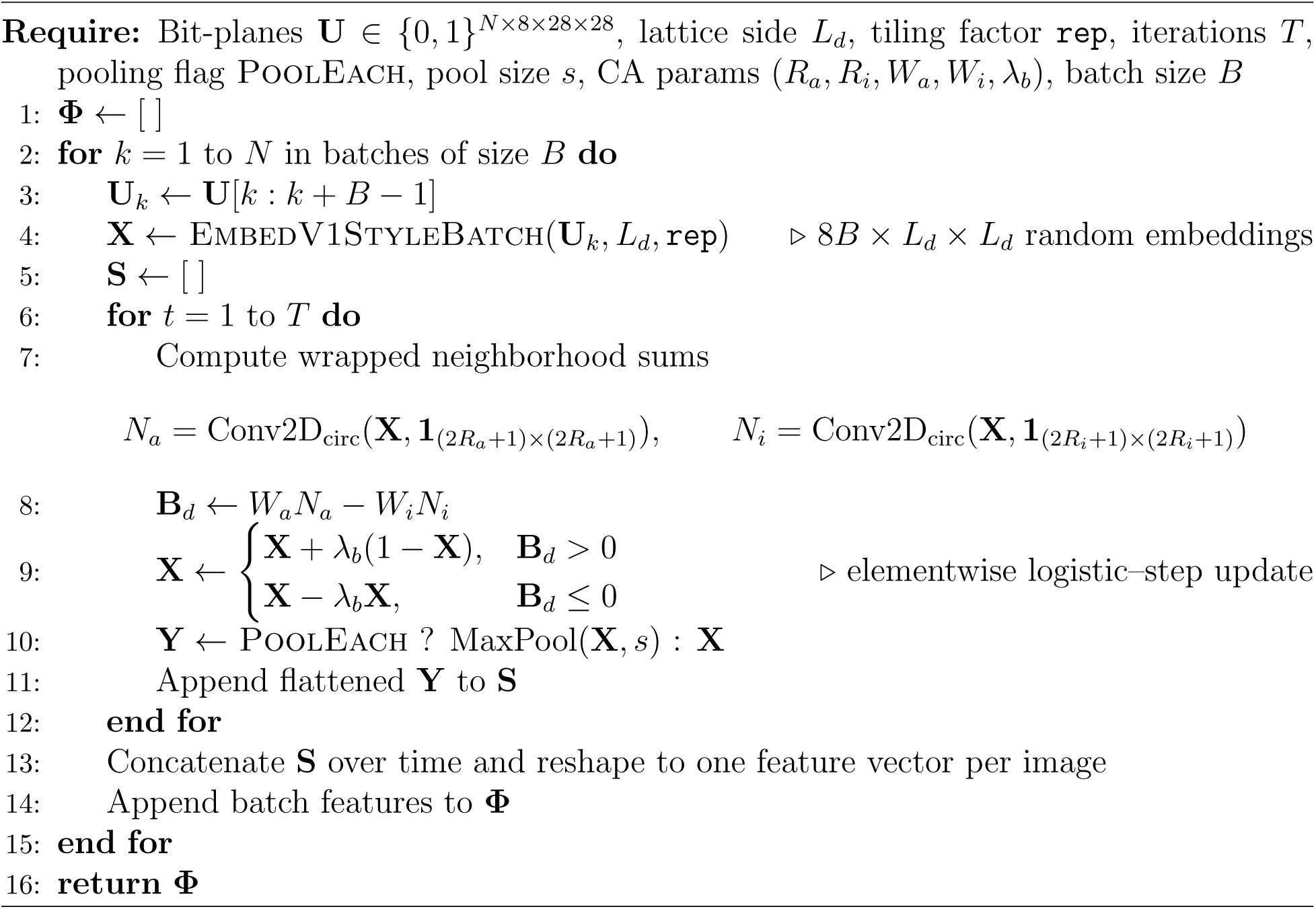

##### Algorithm 11

ConductMNISTExperimentCUDA (current sweep protocol)

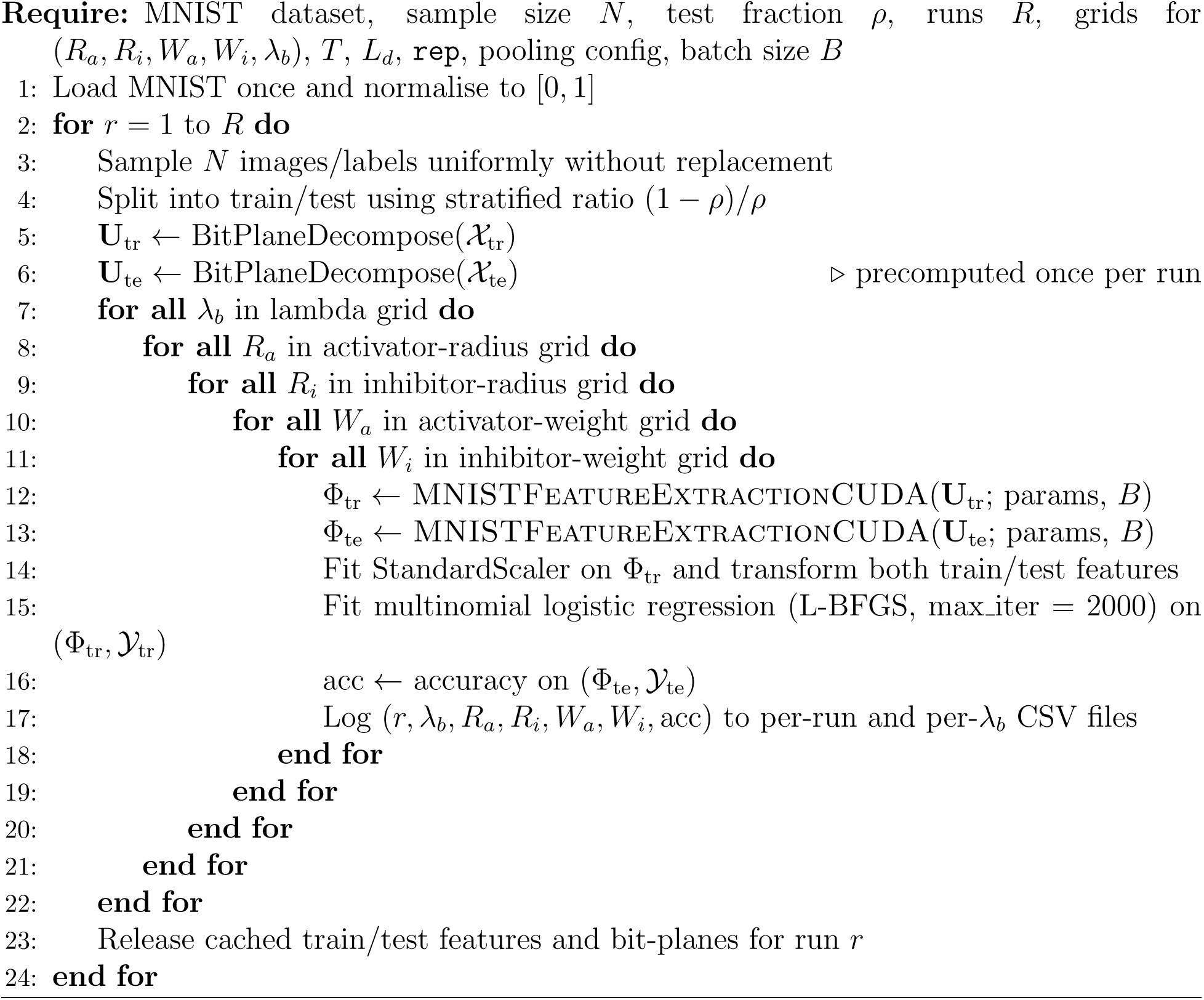

##### Algorithm 12

BestMeanCurveFromSweep

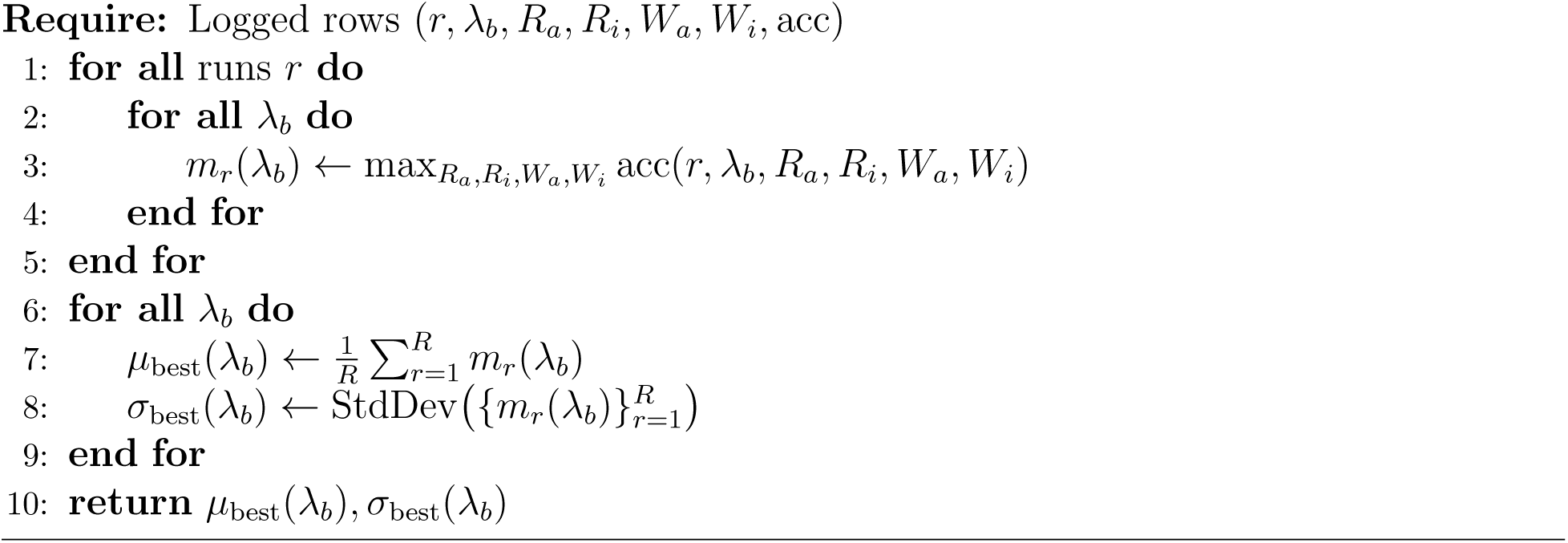

## Data availability

The data generated and analysed during this study are available at https://doi.org/10.5281/zenodo.1945810 Source data underlying the main figures are provided with this paper.

## Code availability

The code used to generate the results in this study is available at https://doi.org/10.5281/zenodo.19458106.

The repository includes the CA simulation code, reservoir-computing pipelines, analysis scripts, and figure-generation scripts.

## Acknowledgements

The authors thank the open-source software and dataset communities whose resources supported the computational analyses.

## Author contributions

J.R. conceived the study, developed the methodology, implemented the code, performed the simulations and analyses, and wrote the manuscript. C.P.B. contributed conceptual input on Turing partial differential-equation models and reviewed and edited the manuscript. A.Z. contributed conceptual input on the reservoir-computing components of the study and reviewed and edited the manuscript. All authors discussed the results and approved the final manuscript.

## Ethics declarations

Not applicable; this study did not involve human participants, human data, animals, clinical samples, or experiments requiring ethical approval.

## Additional information

Correspondence and requests for materials should be addressed to A.Z. The Zenodo record at DOI 10.5281/zenodo.19458106 was verified as public and open at the time of preparing this npj Unconventional Computing submission.

